# Reliability and signal comparison of OPM-MEG, fMRI & iEEG in a repeated movie viewing paradigm

**DOI:** 10.1101/2025.07.14.664811

**Authors:** Olivia R. Christiano, Sebastian Michelmann

## Abstract

Optically pumped magnetometers (OPMs) offer a promising advancement in noninvasive neuroimaging via magnetoencephalography (MEG), but establishing their reliability and comparability to existing methods remains an ongoing endeavor. Here, we evaluated OPM recordings by assessing their test-retest reliability and comparing them to functional magnetic resonance imaging (fMRI) and intracranial electroencephalography (iEEG) recordings. Data were collected from three independent participant groups during repeated viewings of a movie segment. In 7 canonical frequency bands (*δ*: 0.5–4 Hz, *θ*: 4–8 Hz, *α*: 8–12 Hz, *β*: 12–28 Hz, *γ*_1_: 28–46 Hz, *γ*_2_: 55–70 Hz, and HF: 64–116 Hz), as well as a broadband (BB: 0.5–116 Hz) signal, we quantified the signal consistency (1) within individuals, (2) across subjects, and (3) across modalities. OPM exhibited widespread reliability, particularly in lower frequency bands; spatial patterns resembled those of fMRI and iEEG in visual and auditory regions. Cross-modal analyses revealed robust correspondence between OPM and both fMRI and iEEG, including inverse correlations at low frequencies and positive correlations at higher frequencies in the OPM-fMRI comparison, consistent with known relationships between oscillatory power and BOLD responses. Comparisons of signal-to-noise (SNR) estimates further revealed that in some regions, the SNR of cross-modal alignment exceeded within-modality reliability, suggesting that bridging between modalities can sometimes enhance SNR by attenuating reliably shared noise. Our findings demonstrate that OPM consistently captures stimulus-locked neural dynamics that converge with established modalities.

## 1. INTRODUCTION

Advancing our understanding of the neural activity underlying human cognition depends on tools that reliably capture neurophysiological signals with high temporal precision. Optically pumped magnetometers (OPMs) represent a promising innovation in magnetoencephalography (MEG), a noninvasive technique with high temporal resolution (milliseconds) that can (under favorable conditions) measure cortical sources with up to 2– 3 mm precision (Brookes et al., 2021; D. Cohen, 1968, 1972; Hämäläinen et al., 1993; Proudfoot et al., 2014). OPMs address many of the practical limitations associated with superconducting quantum interference devices (SQUIDs) used in conventional MEG systems, whose cryogenic requirements impose substantial operational costs and technical constraints (Andersen et al., 2017; Boto et al., 2017; D. Cohen, 1972; Hämäläinen et al., 1993; Hill et al., 2020): Unlike SQUIDs, OPMs operate at room temperature, eliminating the need for cryogenic cooling and the associated costs (Boto et al., 2017). Their lightweight, compact design allows for direct placement on the scalp, which minimizes the sensor-to-brain distance and may consequently enhance signal amplitude and spatial resolution (Hill et al., 2020; Iivanainen et al., 2017; Nugent et al., 2022). Additionally, OPMs can be configured into wearable, adjustable helmets, accommodating naturalistic movement during data acquisition, and enabling greater adaptability to different head sizes and shapes (Boto et al., 2018). These features not only broaden MEG’s accessibility and versatility in research but also have the potential to improve signal quality by ensuring consistent and close sensor-to-scalp proximity across individuals (Boto et al., 2016, 2017, 2018; Brookes et al., 2021; Hill et al., 2020; Nugent et al., 2022; Rea et al., 2022).

OPM–MEG (hereafter referred to as OPM) is consequently positioned to be a major advancement in neuroimaging, yet its utility hinges on validating whether it captures reliable and meaningful neural signals. While it has been shown that OPMs can detect canonical MEG signals such as evoked responses, oscillatory activity, functional localization, and large-scale connectivity (An et al., 2022; Boto et al., 2021; Rier et al., 2023; Safar et al., 2024; Sun et al., 2024; Tierney et al., 2018; Wang et al., 2024), their reliability has not been as thoroughly established as the reliability of SQUID based MEG (e.g., Lankinen et al., 2014; Lew et al., 2021; Liu et al., 2022; Martín-Buro et al., 2016).

Moreover, while OPMs have demonstrated strong concordance with SQUIDs across sensor- and source-level analyses, sometimes achieving comparable or greater signal quality with fewer sensors (Iivanainen et al., 2023; Marhl et al., 2022; Rhodes et al., 2023), comparative evidence with other neuroimaging and electrophysiological modalities remains limited. Because different measurement techniques are sensitive to distinct neural processes and noise sources, direct comparisons are needed to clarify both shared and modality-specific components. For instance, EEG and iEEG primarily detect radially oriented electrical currents in cortical gyri, while MEG is most sensitive to tangentially oriented sources located in the sulci. Furthermore, MEG is particularly sensitive in regions closer to the sensors because magnetic field strength diminishes rapidly as it travels away from the source (Hämäläinen et al., 1993). In contrast, fMRI measures vascular responses associated with neural activity, capturing slow hemodynamic changes rather than direct electrophysiological signals. As a novel technique, it is important to understand which neurophysiological signals OPM captures, and critically, to determine whether it does so reliably across individuals and in alignment with signals measured by other established neuroimaging methods.

Here, we systematically compare neural activity recorded with OPM, fMRI, and iEEG during repeated viewings of a continuous audio-visual movie stimulus. We *(i)* quantify OPM signal reliability within individuals, *(ii)* quantify shared OPM signal between individuals, *(iii)* determine the extent to which OPM captures the same stimulus-locked neural activity as fMRI and iEEG, and *(iv)* characterize how its signal reliability varies across these comparisons. To answer these questions, we compute test–retest reliability within individual participants, between different subjects (Intersubject correlations, ISC; Hasson et al., 2004), and between recording modalities. (See Figure 1 for an overview of our analysis framework.) Finally, we compare signal-to-noise ratios (SNR) within OPM and between OPM and fMRI to characterize their variation across levels of comparison.

**Figure 1:**
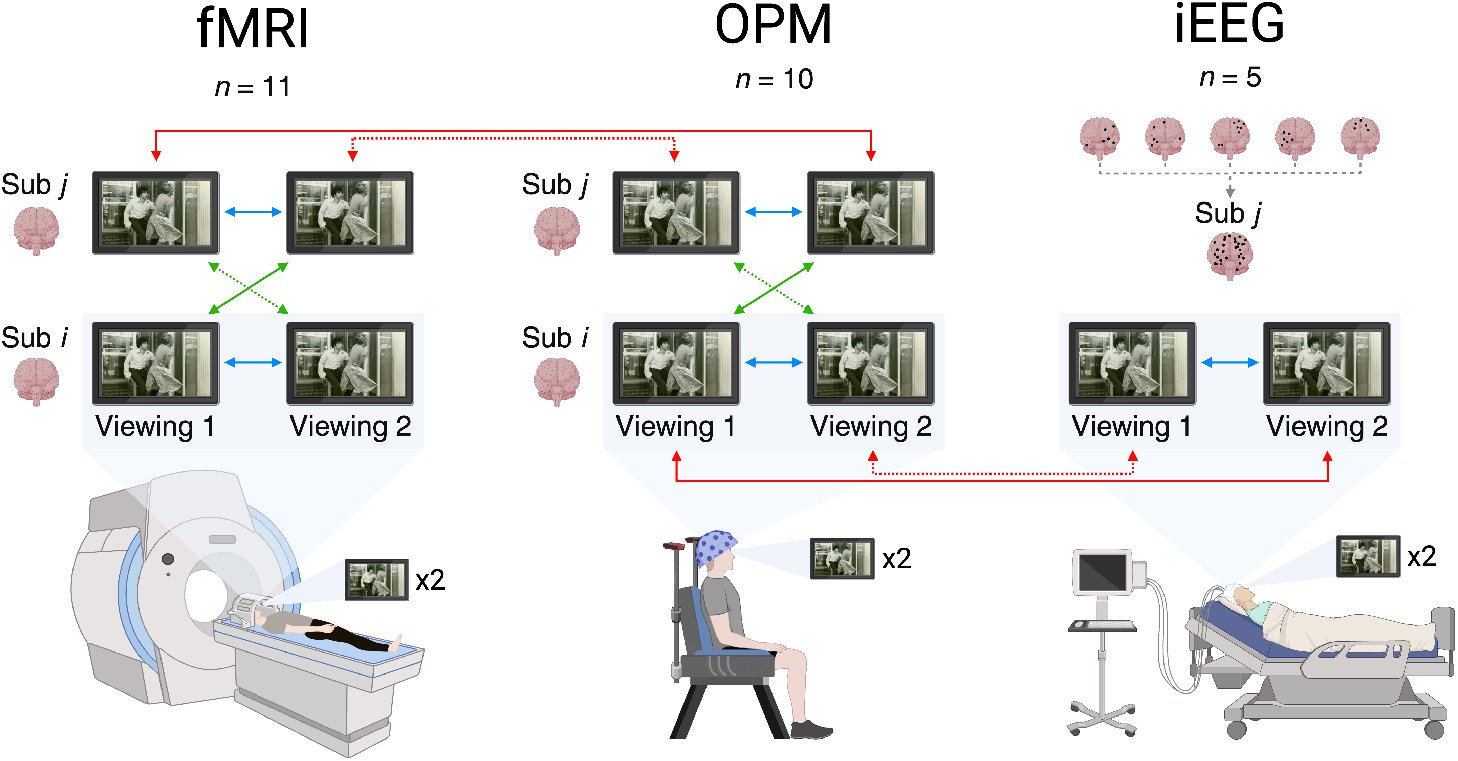
Schematic of correlation analyses. Within-subject correlations (blue) were computed between the first and second viewing for each subject and averaged across the respective cohort; for iEEG, electrodes were treated as a single subject. Between-subject correlations (green) were computed across viewings, averaging correlations from viewing 1 to viewing 2 (solid) and vice versa (dashed), comparing each subject to the rest of their cohort. Between-method correlations (red) were computed similarly, comparing each subject *i* in one modality to each subject *j* in another, or to the single-subject iEEG dataset; fMRI to iEEG comparisons were conducted but are not depicted here for simplicity. Within-viewing analyses were also conducted and are reported in the Supplementary Materials. Notes: Movie was presented in color. Diagrams created in https://BioRender.com/epg2wkb. Image used in this schematic: Warner Bros. (1974). [Al Pacino and Penelope Allen in Dog Day Afternoon] [Photograph]. Wikimedia Commons.

**Figure 2:**
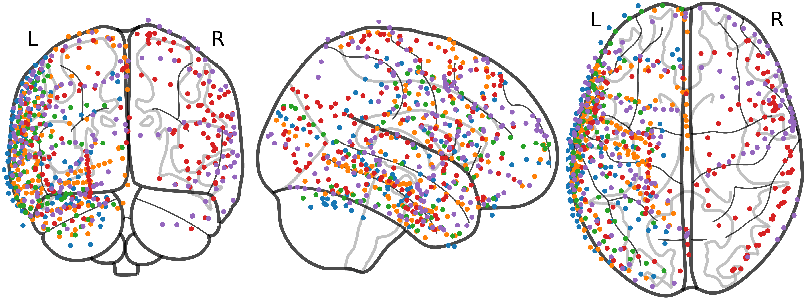
Location of iEEG electrodes on cortical surface. iEEG electrode placements are color-coded by subject: Subject 1 (red), Subject 2 (green), Subject 3 (blue), Subject 4 (orange), Subject 5 (purple).

## 2. METHODS

Neural activity was recorded from three independent subject groups as they watched the same segment of the feature film Dog Day Afternoon (DDA; Lumet, 1975) twice. The datasets used in this study comprised recordings from ten OPM subjects (Rier et al., 2023), eleven fMRI subjects (Haufe et al., 2018), and five iEEG subjects (Honey et al., 2012). Comprehensive details on study subjects, data acquisition, experimental settings, and preprocessing steps can be found in the corresponding articles; however, a succinct description of each study’s methodology is provided below. Unless otherwise specified, all data were processed using custom MATLAB scripts and the FieldTrip toolbox (Oostenveld et al., 2010).

### 2.1 OPM (Rier et al., 2023)

#### 2.1.1 Subjects and experimental settings

OPM data were originally collected at the University of Nottingham as part of a separate study (Rier et al., 2023). The OPM cohort consisted of ten subjects (4 female, 6 male, all right-handed; age: *M* = 31 years, SD = 8) who provided written informed consent to participate in the study. The study protocol was approved by the University of Nottingham Medical School Research Ethics Committee.

Subjects were presented with a 600 s clip of DDA twice, with approximately 1–2 minutes between each viewing. The movie clips were presented at a visual angle of ∼13° horizontally and ∼9° vertically, approximately 80 cm from the subjects’ heads, and audio was delivered via a waveguide connected to speakers outside of the scanning room (Rier et al., 2023).

#### 2.1.2 Signal acquisition

OPM data were recorded using triaxial OPM sensors (QuSpin, Inc., Colorado, USA) mounted in a 3D-printed helmet (Cerca Magnetics Ltd., Nottingham, UK), selected to best fit each participant. Each sensor has a noise floor of ∼13 fT/sqrt (Hz), a bandwidth of ∼150 Hz, and provides three independent channels. Sensors were distributed evenly across the scalp, enabling approximately whole-head coverage, which resulted in an average of ∼168 channels per subject (range: 165–174). Recordings took place in an OPM-optimized magnetically shielded room (Cerca Magnetics Ltd., Nottingham, UK) equipped with background magnetic field control via reference OPMs, biplanar electromagnetic coils, and a motion tracking camera system. Once magnetic field stabilization procedures were complete, data were recorded as participants viewed the movie segment (600 s per run, sampled at 1200 Hz). Co-registration of sensors to anatomy was performed by aligning 3D digitizations of participants’ heads, collected with and without the helmet on, to MRI-derived surfaces (Rier et al., 2023).

#### 2.1.3 Preprocessing

OPM data were minimally preprocessed. Channels were excluded based on the initial preprocessing performed by Rier et al., 2023, which removed sensors with near-zero variance, resulting in an average of 164 usable channels per subject (range: 161– 166). To minimize edge artifacts, MEG data were padded with 1 s before and after the 600 s movie segment, then bandpass filtered (0.5–150 Hz), notch filtered at 50, 100, and 120 Hz, and down-sampled to 256 Hz.

#### 2.1.4 Source reconstruction

Source reconstruction was performed using linear constrained minimum variance (LCMV) beamforming (Veen et al., 1997). Individual T1-weighted MRIs were segmented and used to construct single-shell head models (Nolte, 2003), co-registered with subject-specific sensor locations. An 8 mm-spaced template grid in MNI space was nonlinearly warped to each participant’s anatomy to obtain individualized source positions for corresponding MNI coordinates. Sources were estimated using the data covariance computed separately for each movie segment across the 600 s time window, with 5% regularization applied to the covariance matrix to stabilize the beamformer solution. The dipole orientation was fixed to the direction of maximum amplitude. Separate filters were computed for each movie repetition in each of the 10 participants.

For comparison to iEEG, source reconstruction of the OPM data was repeated using a shared grid defined by the MNI coordinates of 656 iEEG electrodes. These coordinates were nonlinearly warped to each participant’s anatomical space, and source activity was reconstructed directly at these locations using the same preprocessed MEG data, head models, and beamforming procedure described above.

Because source reconstruction parameters can influence the spatial distribution of reconstructed activity, and no single approach is uniquely correct, we performed additional analyses to demonstrate the robustness of our findings. Specifically, we recomputed all main OPM analyses using a minimum-norm-estimation (MNE) solution that did not consider the data covariance structure or include regularization. These results, reported in the Supplementary Materials, show quantitatively consistent reliability estimates, albeit with broader spatial extent, demonstrating that the effects reported here are not driven by beamformer-specific assumptions.

#### 2.1.5 Amplitude of the analytic signal

To extract frequency-specific activity, the source-reconstructed OPM data were bandpass filtered according to the following frequency bands: *δ* (0.5–4 Hz), *θ* (4–8 Hz), *α* (8–12 Hz), *β* (12–28 Hz), two ranges within the *γ* band (*γ*_1_: 28–46 Hz, *γ*_2_: 55–70 Hz), a high frequency band (HF: 64–116 Hz), and a broad band (BB: 0.5–116 Hz), using a 4th order Butterworth IIR filter. We then estimated the log-transformed amplitude at each source location by taking the logarithm of the absolute value of the analytic signal obtained via the Hilbert transform, following standard recommendations for oscillatory amplitude analysis (M. X. Cohen, 2014). This computation was performed separately for each subject, movie viewing, and frequency band.

#### 2.1.6 Cross-modal temporal alignment

OPM data were trimmed to match the adjusted 297 s time axis of the fMRI data (see Section 2.2) by removing the first 16.74 s and the final 286.26 s of the original 600 s recording. Time courses of log-transformed amplitude were resampled to 20 Hz to enable subsequent cross-modal analyses and ensure temporal alignment across modalities.

### 2.2 fMRI (Haufe et al., 2018)

#### 2.2.1 Subjects and experimental settings

fMRI data were originally collected at Princeton University for the purpose of another study (Haufe et al., 2018). The fMRI sample consisted of eleven subjects (6 female, 5 male; age range: 20-35 years) who provided written informed consent to participate in the study, approved by the Princeton University Institutional Review Board. All subjects were fluent in English, with normal or corrected-to-normal vision, and no history of psychiatric or neurological disorders.

Subjects viewed a 325 s clip from DDA twice, with approximately 11 minutes between repetitions, as part of a larger study. Movie clips were displayed (subtended 20° horizontally and 16° vertically) using the Psychophysics Toolbox in MATLAB, synchronized with MRI data acquisition, and presented with audio delivered via in-ear headphones (Haufe et al., 2018).

#### 2.2.2 Signal acquisition

MRI data were collected using a 3T Skyra scanner (Siemens, Munich, Germany) with a 16-channel head coil. Functional images were acquired with a gradient-echo EPI sequence (3 *×* 3 *×* 4 mm resolution, 27 interleaved slices, TR = 1.5 s, TE = 30 ms, flip angle = 72^◦^, FOV = 192 *×* 192 mm^2^, GRAPPA iPAT = 2); high-resolution structural scans were obtained in each session using an MPRAGE sequence (0.9375 mm isotropic, TR = 1.9 s, TE = 2.1 ms, flip angle = 9^◦^, FOV = 240 *×* 240 mm^2^) (Haufe et al., 2018).

#### 2.2.3 Spatial registration

Each subject’s data were aligned to MNI space using AFNI’s linear (3dAllineate) and nonlinear (3dQWarp) registration tools (Cox, 1996), resampled to a 4 mm grid using linear interpolation, and spatially smoothed using a 10 mm full-width-at-half-maximum Gaussian kernel, resulting in the retained volumetric data (32,798 voxels; Haufe et al., 2018).

#### 2.2.4 Preprocessing

Functional data were preprocessed by the original authors as follows: slice-time- and motion-corrected (AFNI’s 3dvolreg) data were linearly detrended, and the discrete Fourier transform (and its inverse) were used to apply a high-pass filter at 0.01 Hz. To ensure signal stability and exclude the blank screen period at the conclusion of the recording, the first 15 s and final 13 s were removed, yielding 297 s of data (Haufe et al., 2018).

#### 2.2.5 Cross-modal alignment

To align the fMRI data to the OPM source grid, we interpolated fMRI activity from MNI-space coordinates onto the template source model used in the OPM beamforming analysis. For each grid point inside the brain volume, we identified the nearest fMRI coordinate(s) based on the minimum Euclidean distance and averaged the time series of all voxels assigned to each corresponding grid point. This yielded spatially aligned fMRI data on the same cortical grid used for OPM source projection.

For analyses comparing fMRI and iEEG, we aligned the fMRI data to the MNI-space coordinates of the 656 iEEG electrodes. For each electrode, we identified all fMRI voxels within a 6 mm radius based on Euclidean distance and averaged their time series. This yielded a version of the fMRI dataset spatially down-sampled to the iEEG grid, allowing direct comparison across modalities at matched anatomical locations. Voxels outside this radius were excluded from the average, and voxel counts per electrode were tracked to normalize the resulting time series.

To ensure temporal alignment across imaging modalities, the fMRI time axis was adjusted to account for hemodynamic delay. The signal was shifted by the lag that maximized the cross-correlation between the amplitude envelope of the movie audio and the time series of the 20 locations nearest to an anatomical mask of auditory cortex regions defined by the AAL atlas (Tzourio-Mazoyer et al., 2002). Auditory cortex was selected as the reference region because its BOLD response has been shown to reliably track continuous auditory input during naturalistic movie viewing, providing a robust, stimulus-locked signal for temporal alignment (Hasson et al., 2010; Mukamel et al., 2005). Preprocessed data were then linearly interpolated to a resolution of 20 Hz.

### 2.3 iEEG (Honey et al., 2012)

#### 2.3.1 Subjects and experimental settings

iEEG data were collected at New York University for the purpose of a separate study (Honey et al., 2012). The iEEG cohort consisted of five subjects (4 female, 1 male; age range: 20–47 years) from the Comprehensive Epilepsy Center of the New York University School of Medicine experiencing pharmacologically refractory complex partial seizures. All patients provided informed consent both before and after electrode implantation in accordance with National Institutes of Health guidelines administered by the local Institutional Review Board.

Patients viewed the same intact 325 s clip of DDA twice, each followed by a 30 s fixation period, with approximately 12 minutes between intact repetitions. Stimuli presented using PsychToolbox Extensions for MATLAB were displayed approximately 40–60 cm from the subjects’ eyes, on a bedside laptop.

#### 2.3.2 Signal acquisition

Recordings were obtained from 922 subdural electrodes in implanted arrays (8 *×* 8, 4 *×* 8, or 1 *×* 8 configurations) spaced 10 mm apart and 2.3 mm in exposed diameter; signals were referenced to skull screws and acquired at 30 kHz using a custom system built on the NSpike framework (L.M. Frank and J. MacArthur, Harvard University Instrument Design Laboratory, Cambridge, MA), which incorporates a 0.6 Hz hardware high-pass filter (Honey et al., 2012).

#### 2.3.3 Spatial registration

Electrode positions were determined using a combination of T1 images (collected pre- and post-implantation) labeling, intraoperative photos, and a MATLAB tool incorporating known electrode geometry (Yang et al., 2012)); SPM’s DARTEL algorithm (Ashburner, 2007) was then used to warp each subject’s anatomy and individual electrode coordinates to MNI space (Honey et al., 2012). Note that, unlike Haufe et al. (2018) and Honey et al. (2012), we did not exclude electrodes based on hemisphere location; thus, all 656 artifact-free, labeled, and MNI-registered electrodes were retained. Given the availability of MNI coordinates as precise estimates of contact locations, we analyzed iEEG data in MNI space rather than estimating anatomical sources.

#### 2.3.4 Preprocessing

Data from all five subjects were first aggregated into a single dataset. Signals were high-pass (0.5 Hz), low-pass (150 Hz), and notch filtered at 60 and 120 Hz, following (Honey et al., 2012), using 4th-order Butterworth IIR filters implemented with FieldTrip. The resulting data were then down-sampled to 256 Hz.

#### 2.3.5 Amplitude of the analytic signal

To estimate the time-resolved amplitude of iEEG activity, broadband signals were bandpass filtered into the same frequency bands as the OPM data using 4th-order Butterworth IIR filters, and the log-transformed amplitude was computed as described in Section 2.1.5.

#### 2.3.6 Cross-modal temporal alignment

For the 338 s of iEEG data, we validated the temporal alignment with the stimulus based on the lag that maximized the cross-correlation between the envelope of the movie’s audio-trace and auditory responses in the high-frequency band (0.5– 116 Hz). This signal was selected because it is a well-established signature of temporally precise, stimulus-locked auditory engagement (Nourski et al., 2009; Sinai et al., 2009), and alignment was further confirmed by visual inspection of peaks in the cross-correlogram across frequency bands. To this end, high-frequency activity from the aggregated dataset was averaged across the 20 locations nearest to an anatomical mask of auditory cortex regions defined by the AAL atlas (i.e., we followed the same procedure used to temporally align the fMRI to the movie). Data were then trimmed to the same 297 s as the OPM and fMRI. Time courses of log-transformed amplitude were, again, resampled to 20 Hz.

### 2.4 Analyses

#### 2.4.1 Overview and methodological considerations

To identify the shared neural responses elicited across repeated movie viewings, we evaluated the consistency of activity within individuals, between individuals, and across recording modalities. All analyses were performed on the data segments temporally aligned to the movie stimulus and trimmed to identical duration across modalities. All correlation analyses were performed within separate frequency bands; for fMRI analyses the BOLD signal was used. For OPM and fMRI, coefficients were computed using Pearson product-moment correlations, whereas for iEEG, Spearman rank correlations were used to account for potential outliers in signal amplitudes due to epileptic activity. Because iEEG electrode locations varied across subjects, between-subject comparisons could not be performed within the iEEG dataset.

#### 2.4.2 Within-subject comparisons

For each of the three modalities, subject-level empirical correlations were computed between the time series from the first and second movie viewings.

#### 2.4.3 Between-subject comparisons

Pairwise correlations were computed bidirectionally between one viewing from each subject (first/second) with the alternate viewings (second/first) of the remaining subjects in the same cohort (OPM or fMRI).

#### 2.4.4 Between-method comparisons

##### 2.4.4.1 OPM to fMRI

Cross-modal consistency between OPM and fMRI was assessed by computing Pearson correlations between each movie viewing (first/second) from each OPM subject and the alternate viewing from each fMRI subject (second/first). Fisher *z*-transformed correlations were then averaged across OPM and fMRI subjects.

##### 2.4.4.2 OPM to iEEG

Spearman correlations were computed between each viewing from each OPM subject (first/second viewing) and the alternate viewing from the single-subject iEEG dataset (second/first viewing). Fisher *z*-transformed correlation coefficients were averaged across OPM subjects.

##### 2.4.4.3 fMRI to iEEG

To comprehensively compare all three imaging modalities, we evaluated the consistency between iEEG and fMRI to reproduce the findings of Haufe et al. (2018) with our analysis approach. Spearman correlations were computed bidirectionally between one viewing from each fMRI subject and the alternate viewing from the single-subject iEEG dataset and averaged across fMRI subjects.

#### 2.4.5 Statistical significance and analysis framework

Correlation estimates are sensitive to the autocorrelational structure of neural time series (Schaworonkow et al., 2015; Woolrich et al., 2001), which may be magnified by temporal down-sampling and smoothing (James et al., 2019; Mejia et al., 2015; Pajula & Tohka, 2014; Zarahn et al., 1997). To account for these dependencies, we assessed statistical significance using standardized metrics derived from modality-specific null distributions

(Haufe et al., 2018). Unlike correlation coefficients, which quantify the magnitude of association between two signals, *z*-scores reflect how extreme an observed correlation is relative to the expected distribution under the null, given the temporal structure and noise characteristics of the data. As a result, modest correlation values may yield large *z*-scores if the null distribution is narrow, whereas larger correlation coefficients may not translate into large *z*-scores when the null distribution is broader. We further illustrate the influence of temporal resolution on correlation estimates by computing Pearson correlations of OPM and iEEG at the same sampling rate as fMRI. To this end, we first downsampled the OPM and iEEG data to 0.67 Hz (the temporal frequency of the 1.5s TR in fMRI) and then repeated our within-subject correlation analysis (see Table S1). Crucially, because we aimed to quantify signal consistency rather than to maximize correlation coefficients, we resampled all estimates to 20 Hz, a resolution commonly used in the analysis of power spectra in human electrophysiology (e.g., Michelmann et al., 2016; see also: M. X. Cohen, 2014). We then computed surrogate null distributions using the iterative amplitude-adjusted Fourier transform (iAAFT; Schreiber and Schmitz, 1996), and standardized empirical correlations by deriving *z*-scores relative to the null distribution. We further assessed the significance of observed correlations based on the distribution of averages of surrogate correlation. Specifically, we performed the following steps:

1. Empirical correlations were computed from pairs of real-time series at a given channel (i.e., source position, voxel, or electrode) in a comparison of interest (between viewings, subjects, or modalities). Subject-level empirical correlations were then Fisher *z*-transformed and averaged across subjects, producing a grand-averaged (GA) empirical correlation coefficient for each channel.
2. We generated 100 surrogate datasets for each time series (movie viewing, subject, frequency band, and modality) and computed surrogate correlations, where one of the empirical time series in a given comparison was replaced with a phase-randomized surrogate time series. The 100 surrogate correlations that correspond to an empirical correlation were also normalized using the Fisher *z*-transform.
3. Null distributions of averages were then bootstrapped by randomly sampling one surrogate correlation per subject and averaging across subjects to form a single set of GA surrogate correlation coefficients analogous to the empirical GA correlations. This averaging was repeated 10,000 times per comparison, randomly resampling surrogates (i.e., selecting 1 out of 100 options per subject for each surrogate GA). This yielded a null distribution of 10,000 GA surrogate correlation coefficients per channel (Stelzer et al., 2013)
  3a. For the iEEG within-subject analysis, subject-level bootstrapping could not be performed. Instead, we repeated step 2 with 1,000 surrogate datasets per channel, using the resulting surrogate correlations directly as the null distribution.
4. *p*-values were calculated for each channel as the proportion of the surrogate GA correlations that exceeded the empirical GA coefficient. The Benjamini-Hochberg correction was applied to the empirically derived *p*-values across channels in each analysis, effectively controlling the false discovery rate (FDR) at a level of *q* = .05.
5. For visualization and interpretation, the empirical GA correlations at each channel were standardized relative to the channel’s surrogate distributions by computing *z*-scores: 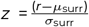, where *µ*_surr_ and *σ*_surr_ denote the *M* and *SD* of the surrogate null distribution. In other words, for each channel we compared a single empirical value, corresponding to the average of subject-level correlations, to a null distribution in which each sample reflects a surrogate value computed as the average of randomly selected subject-level surrogate correlations, thereby preserving the relationship between empirical and surrogate estimates. These *z*-scores reflect estimates of SNR and enable comparison of SNR on a common scale across modalities and analyses.

#### 2.4.6 Signal-to-noise (SNR) comparisons

To evaluate the relative SNR of OPM signals within and across subjects, we compared the channel-wise SNR estimates by subtracting the within-subject *z*-scores from the between-subject *z*-scores for each frequency band. Positive values in the resulting maps indicate greater consistency of neural responses across individuals than within, whereas negative values indicate greater consistency within than across individuals. Cluster-based thresholding was used to identify contiguous channels exceeding the 97.5th or falling below the 2.5th percentile of the observed SNR distribution. The sum of *z*-scores for each cluster was used to identify the maximum clusters for both the positive and negative tails. We then assessed the significance of the maximum clusters relative to 10,000 surrogate SNR maps. These were computed as the difference between randomly sampled surrogate-within- and between-subject correlation sets derived from the null distributions computed in previous analyses. Empirical *p*-values were calculated as the proportion of surrogate absolute cluster sums that were bigger than the empirical maximum cluster sums. We controlled the false discovery rate at *q* = .025 per frequency band.

We performed the same analysis to compare between-subject correlations in OPM to between-method correlations in OPM and fMRI. For each frequency band, we computed channel-wise SNR maps by subtracting the OPM’s between-subject *z*-scores from the corresponding *z*-scores in the OPM to fMRI comparison. Because cross-modal comparisons may yield meaningful inverse relationships, we used the absolute value of the between-method *z*-scores to represent the magnitude of the correlation, regardless of its direction. Thus, resulting positive values indicate greater consistency between OPM and fMRI than across OPM subjects alone, whereas negative values indicate greater consistency across OPM subjects than between OPM and fMRI. Cluster-based permutation testing and significance evaluation followed the same procedure as described above. It is important to note that clusters reflect regions with significant differences, not areas where reliability was equally strong across comparisons.

In addition to these primary SNR analyses, we conducted supplemental contrasts examining modality-specific and cross-modal reliability patterns, which are reported in the Supplementary Materials.

## 3. RESULTS

### 3.1 Within-subject comparisons

Across all three methods, we observed significant reliability in spatially distinct but functionally meaningful areas. For OPM, reliable signal, indicated by channels surviving correction for multiple comparisons (MC), was observed across all frequency bands and spanned regions involved in visual processing, higher-order associative functions, and motor control (Figure 3A). Peak *z*-scores across frequency bands ranged from 5.28 (*γ*_2_) to 16.04 (*θ*), with the highest proportion of channels surviving FDR correction in the *α* band (82.0%; *M*_*z*_ = 5.26, *SD*_*z*_ = 2.09). Signal reliability was broadly distributed in the lower frequency bands (*δ, θ, α*, and *β*), with maxima localized to the left and right cuneus; in contrast, higher frequency bands (*γ*_1_, *γ*_2_, and HF) displayed more focal distributions, with peaks in the precentral gyrus, medial frontal gyrus, and middle frontal gyrus, respectively (see Table 1, OPM). Interestingly, in the 28–116 Hz range, *z*-scores decreased systematically with increasing frequency, but the spatial distributions of reliable effects followed an inverse pattern. For instance, while the HF band had a lower maximum reliability (peak *z* = 5.28) it also demonstrated the greatest proportion of channels surviving MC correction (26.6%) compared to the *γ*_1_ (peak *z* = 6.01, % MC = 18.4) and *γ*_2_ bands (peak *z* = 5.85, % MC = 8.7). This pattern is consistent with prior work indicating that activity in higher-frequencies is driven by distinct neural processes, with *γ* band activity reflecting narrow band network oscillations and HF activity indicating broader population spiking; as such, *γ* oscillations may be locally strong, but spatially sparse, whereas HF activity can appear more spatially distributed but weak relative to noise (Ray & Maunsell, 2011). Finally, the broadband signal exhibited the highest overall reliability, with peak *z*-scores exceeding those of any individual frequency band and a spatial distribution largely overlapping with low-frequency visual regions (Figure 3A).

**Table 1:** Within-subject reliability statistics across modalities and frequency bands.

| Modality | Frequency | $M_z$ | $SD_z$ | % MC | $r_{\max}$ | $z$ | Maxima | |
| --- | --- | --- | --- | --- | --- | --- | --- | --- |
|  |  |  |  |  |  |  | MNI Coord. | AAL Label (Left/Right) |
| OPM | $\delta$ 0.5–4 Hz | 5.00 | 2.19 | 81.2% | 0.08 | 13.42 | 8, -96, 16 | Cuneus (R) |
| | $\theta$ 4–8 Hz | 4.86 | 2.54 | 77.0% | 0.08 | 16.04 | 8, -96, 16 | Cuneus (R) |
| | $\alpha$ 8–12 Hz | 5.26 | 2.09 | 82.0% | 0.08 | 13.57 | 0, -88, 32 | Cuneus (L) |
| | $\beta$ 12–28 Hz | 4.45 | 1.89 | 79.3% | 0.04 | 12.28 | -8, -96, 24 | Cuneus (L) |
| | $\gamma_1$ 28–46 Hz | 3.15 | 0.67 | 18.4% | 0.02 | 6.01 | -40, -16, 64 | Precentral (L) |
| | $\gamma_2$ 55–70 Hz | 3.31 | 0.64 | 8.7% | 0.02 | 5.85 | 0, 56, 0 | Frontal Sup Medial (L) |
|  | HF 64–116 Hz | 2.96 | 0.59 | 26.6% | 0.02 | 5.28 | 40, 56, 8 | Frontal Mid (R) |
|  | BB 0.5–116 Hz | 7.56 | 3.34 | 94.2% | 0.07 | 20.76 | 0, -96, 24 | Cuneus (L) |
| fMRI | – | 4.41 | 2.44 | 55.5% | 0.35 | 13.75 | -56, -16, 0 | Temporal Mid (L) |
| iEEG | $\delta$ 0.5–4 Hz | 3.63 | 1.59 | 30.9% | 0.29 | 12.35 | -58, -14, 48 | Postcentral (L) |
| | $\theta$ 4–8 Hz | 3.57 | 1.48 | 34.8% | 0.22 | 9.79 | 65, -18, 25 | SupraMarginal (R)<sup>a</sup> |
| | $\alpha$ 8–12 Hz | 3.80 | 1.63 | 23.3% | 0.25 | 10.75 | -68, -11, 8 | Temporal Sup (L) |
| | $\beta$ 12–28 Hz | 3.97 | 1.42 | 17.4% | 0.11 | 9.74 | -67, -21, 16 | SupraMarginal (L) |
| | $\gamma_1$ 28–46 Hz | 3.54 | 1.13 | 11.0% | 0.09 | 7.80 | -48, -54, -24 | Temporal Inf (L) |
| | $\gamma_2$ 55–70 Hz | 4.08 | 1.44 | 11.7% | 0.12 | 8.30 | 65, -20, 15 | Temporal Sup (R) |
|  | HF 64–116 Hz | 5.69 | 3.82 | 26.7% | 0.34 | 21.26 | -67, -21, 16 | SupraMarginal (L) |
|  | BB 0.5–116 Hz | 3.83 | 1.86 | 40.9% | 0.25 | 12.30 | -68, -11, 8 | Temporal Sup (L) |

**Note.** All statistics are reported for data surviving correction for multiple comparisons (MC). For each modality and frequency band, we report the mean and standard deviation of z-scores ( $M_z$ , $SD_z$ ), the percentage of FDR-surviving channels/voxels/electrodes (% MC), the maximum absolute correlation coefficient ( $r_{\max}$ ), the peak z-score, its MNI coordinate, and the corresponding anatomical label. Superscript a denotes that the first label was found for the electrode with the second highest z-score.

**Figure 3:**
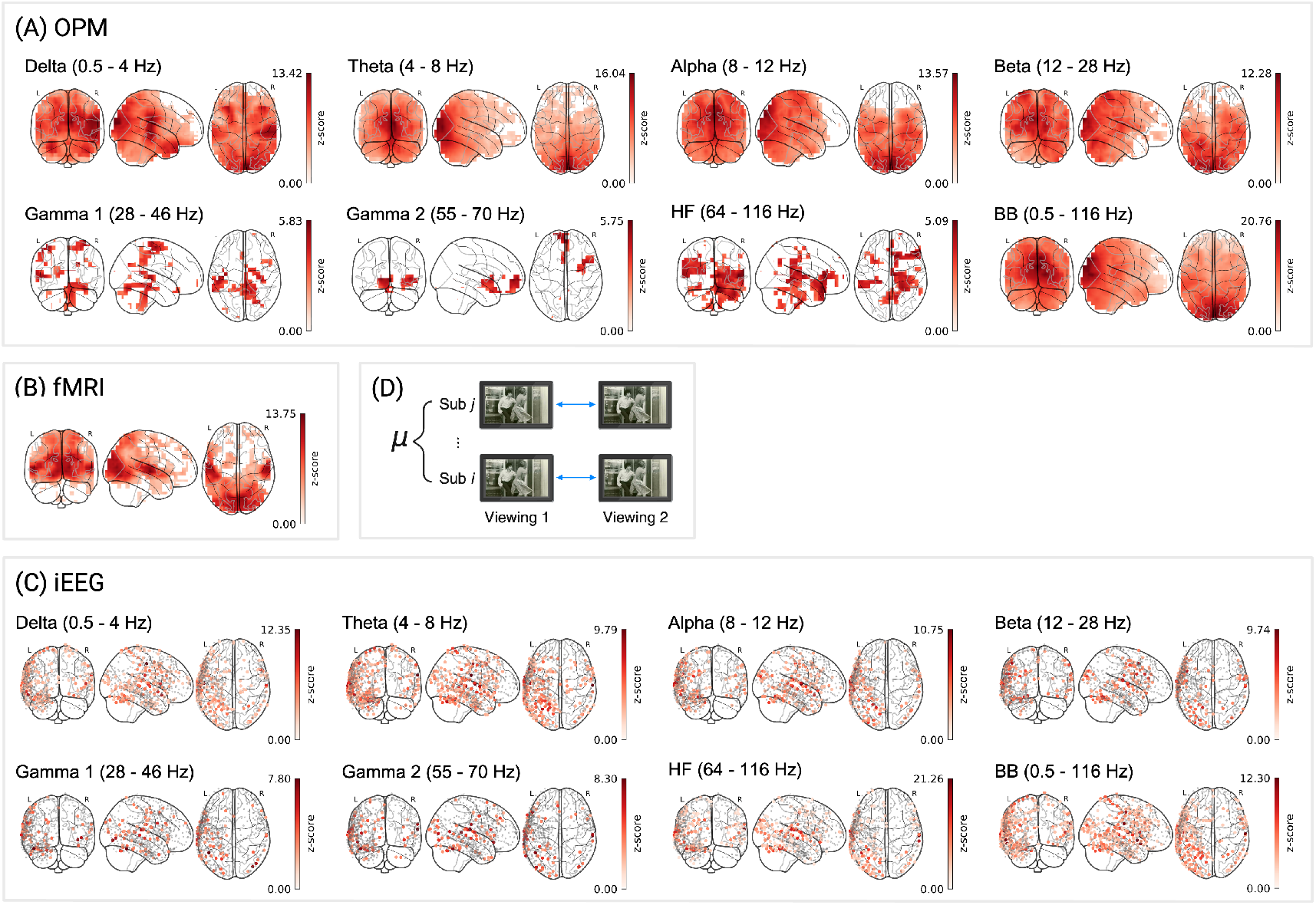
Within-subject reliability for different imaging modalities and frequency bands. FDR-corrected *z*-scores are shown for: (A) OPM, 3,581 source locations across 10 subjects; (B) fMRI, 3,581 voxels across 11 subjects; (C) iEEG, 656 electrodes pooled across five subjects. For iEEG, *z*-scores were computed relative to a null distribution of 1,000 surrogate correlations, without bootstrapping (see Methods). (D) Schematic of within-subject correlation analysis. Data were bandpass filtered to seven frequency bands: *δ* (0.5–4 Hz), *θ* (4–8 Hz), *α* (8–12 Hz), *β* (12–28 Hz), *γ*_1_ (28–46 Hz), *γ*_2_ (55–70 Hz), and HF (64–116 Hz), as well a broadband (0.5–116 Hz). Color scales reflect *z*-scores surviving correction for MC of within-subject correlations, plotted on sensor-specific locations (iEEG) or MNI coordinates (OPM, fMRI).

For fMRI, reliable signal was more spatially localized, with 55.5% of voxels surviving MC correction (Table 1, fMRI). Significant effects were observed primarily in regions associated with auditory and visual processing (see Figure 3B), with a peak *z*-score of 13.75 observed in the left middle temporal gyrus (*M*_*z*_ = 4.41, *SD*_*z*_ = 2.44; see also Table 1, fMRI).

iEEG showed strong within-subject reliability, though spatial extent was more limited, with 11.0–34.8% (see Table 1, iEEG) of electrodes surviving correction for MC across frequency bands. In the lower frequency bands, reliability was strongest in temporal, parietal, and sensorimotor areas (see Figure 3C). Similar to OPM, the broadband iEEG signal yielded the highest proportion of electrodes surviving MC correction (40.9%; Table 1, iEEG) relative to the frequency-decomposed signal; however, the highest FDR corrected *z*-score was observed for the high-frequency band concentrated in the left supramarginal gyrus (*z* = 21.26, *M*_*z*_ = 5.69, *SD*_*z*_ = 3.82).

Although absolute correlation values were highest for fMRI (*r*_max_ = .35) and iEEG (*r*_max_ = .34) compared to OPM (*r*_max_ = .08, compare Table S1), we found that both temporally down-sampling the data and ‘grand averaging’ time series across subjects substantially increased OPM correlations (see Table S1, OPM). When re-computed at 0.67 Hz, matching the temporal resolution of fMRI, OPM peak correlations more than tripled (*r*_max_ = .30), and further increased after grand averaging measurements across subjects before computing correlations (*r*_max_ = .66). These findings illustrate that high test-retest correlations can be achieved with OPM by down-sampling and averaging across subjects. Importantly, estimating the SNR via *z*-scores (see above) confirmed robust statistical reliability.

### 3.2 Between-subject comparisons

OPM exhibited reliable signal across all frequency bands, with effects most pronounced in occipitotemporal and cerebellar regions implicated in perceptual and linguistic processing (see Figure 4A; MacEvoy and Epstein, 2011; Nakatani et al., 2022). Across frequency bands, peak *z*-scores ranged from 5.53 (HF) to 15.29 (*θ*), with the greatest proportion of channels surviving correction for MC in the *β* band (92.8%; *M*_*z*_ = 6.09, *SD*_*z*_ = 2.38; see Table 2, OPM). In lower frequencies, regions of strongest reliability were concentrated in the temporal pole (*δ*), lingual gyrus (*θ*), precuneus (*α*), and superior occipital gyrus (*β*), suggesting pronounced reliability in posterior visual and associative regions. High-frequency bands exhibited more focal effects, with peak values in the cerebellum (*γ*_1_), precuneus (*γ*_2_), and inferior frontal gyrus (HF). Widespread effects were further observed in the braodband (raw) signal, with peak reliability localized to the calcarine fissure (peak *z*-score = 22.13), indicating stable responses across subjects in early visual regions.

**Table 2:**
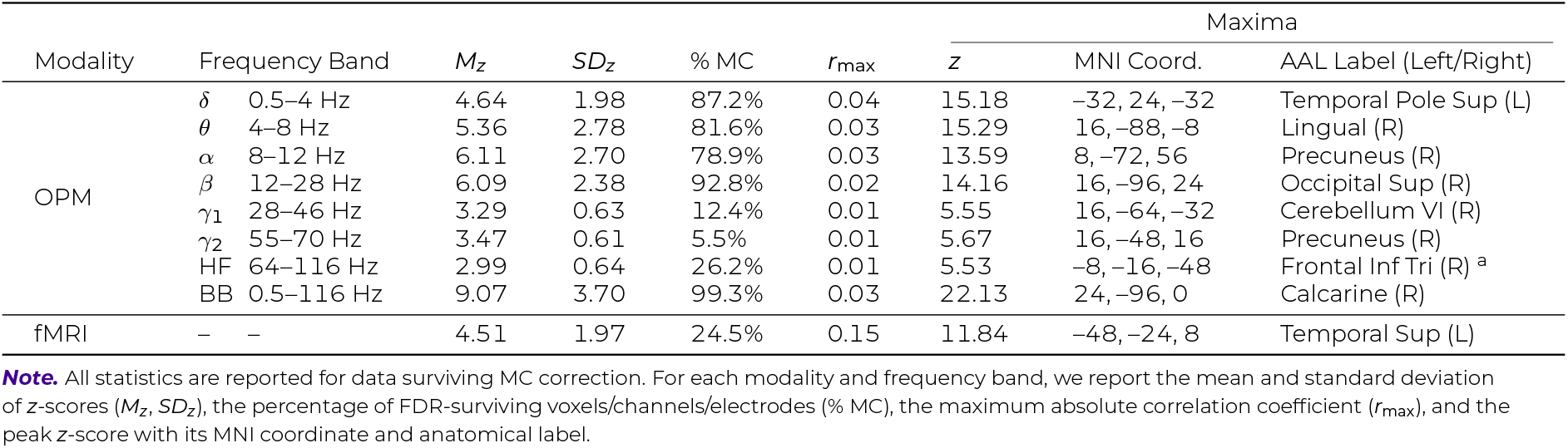
Between-subject reliability statistics for OPM and fMRI.

**Table 3:** Between-method reliability statistics across frequency bands for OPM, fMRI, and iEEG.

| Comparison | Frequency Band | $M_z$ | $SD_z$ | % MC | $r_{\max}$ | Maxima | | |
| --- | --- | --- | --- | --- | --- | --- | --- | --- |
| | | | | | | $z$ | MNI Coord. | AAL Label (Left/Right) |
| OPM to fMRI | $\delta$ 0.5–4 Hz | -2.30 | 3.62 | 5.5% | -0.02 | -8.05 | -24, -56, 56 | Parietal Sup (L) |
| | $\theta$ 4–8 Hz | -3.19 | 3.32 | 9.5% | -0.02 | -9.51 | -24, -56, 72 | Parietal Sup (L) |
| | $\alpha$ 8–12 Hz | -4.37 | 3.05 | 17.8% | -0.03 | -12.42 | -40, -88, 8 | Occipital Mid (L) |
| | $\beta$ 12–28 Hz | -4.02 | 3.14 | 16.1% | -0.02 | -12.00 | -40, -80, 16 | Occipital Mid (L) |
| | $\gamma_1$ 28–46 Hz | 3.74 | 0.00 | 0.0% | 0.01 | 3.74 | 24, -8, 64 | Frontal Sup (R) |
| | $\gamma_2$ 55–70 Hz | 3.45 | 2.13 | 0.7% | 0.01 | 4.72 | 16, -80, 8 | Calcarine (R) |
|  | HF 64–116 Hz | 3.74 | 1.07 | 2.0% | -0.01 | 5.19 | 0, -88, 16 | Cuneus (L) |
|  | BB 0.5–116 Hz | -4.01 | 3.48 | 15.1% | -0.02 | -12.73 | -16, -88, 32 | Occipital Sup (L) |
| OPM to iEEG | $\delta$ 0.5–4 Hz | 3.32 | 1.15 | 6.6% | 0.03 | 4.74 | -37, -45, 69 | Parietal Sup (L) |
| | $\theta$ 4–8 Hz | 3.68 | 1.23 | 9.9% | 0.04 | 7.10 | -21, -55, 78 | Parietal Sup (L) |
| | $\alpha$ 8–12 Hz | 3.57 | 1.89 | 16.3% | 0.05 | 8.11 | -72, -38, 9 | Temporal Mid (L) |
| | $\beta$ 12–28 Hz | 3.55 | 1.09 | 7.9% | 0.02 | 5.68 | -54, -75, 3 | Occipital Mid (L) |
| | $\gamma_1$ 28–46 Hz | – | – | – | – | – | – | – |
| | $\gamma_2$ 55–70 Hz | – | – | – | – | – | – | – |
|  | HF 64–116 Hz | – | – | – | – | – | – | – |
|  | BB 0.5–116 Hz | 3.71 | 1.62 | 24.2% | 0.03 | 8.16 | -29, -49, 73 | Parietal Sup (L) |
| fMRI to iEEG | $\delta$ 0.5–4 Hz | -4.68 | 1.27 | 3.8% | -0.09 | -7.40 | -11, -64, 75 | Precuneus (L) |
| | $\theta$ 4–8 Hz | -3.76 | 2.12 | 7.6% | -0.08 | -7.09 | -74, -28, 3 | Temporal Mid (L) |
| | $\alpha$ 8–12 Hz | -4.37 | 1.20 | 6.1% | -0.08 | -7.96 | -46, -88, 17 | Occipital Mid (L) |
| | $\beta$ 12–28 Hz | -4.11 | 1.13 | 4.1% | -0.05 | -7.48 | -35, -91, 22 | Occipital Mid (L) |
| | $\gamma_1$ 28–46 Hz | 3.77 | 1.72 | 4.0% | 0.03 | 5.96 | -72, -38, 9 | Temporal Mid (L) |
| | $\gamma_2$ 55–70 Hz | 4.53 | 1.26 | 3.5% | 0.05 | 7.54 | -21, -55, 78 | Parietal Sup (L) |
|  | HF 64–116 Hz | 4.97 | 1.44 | 5.9% | 0.07 | 7.71 | -46, -83, 7 | Occipital Mid (L) |
|  | BB 0.5–116 Hz | -4.40 | 1.30 | 8.5% | -0.08 | -7.57 | -46, -88, 17 | Occipital Mid (L) |

**Note.** All statistics are reported for data surviving correction for MC. For each comparison and frequency band, we report the mean and standard deviation of $z$ -scores ( $M_z$ , $SD_z$ ), the percentage of voxels/electrodes surviving correction (% MC), the maximum absolute correlation coefficient ( $r_{\max}$ ), and the peak $z$ -score with its MNI coordinate and anatomical label.

**Figure 4:**
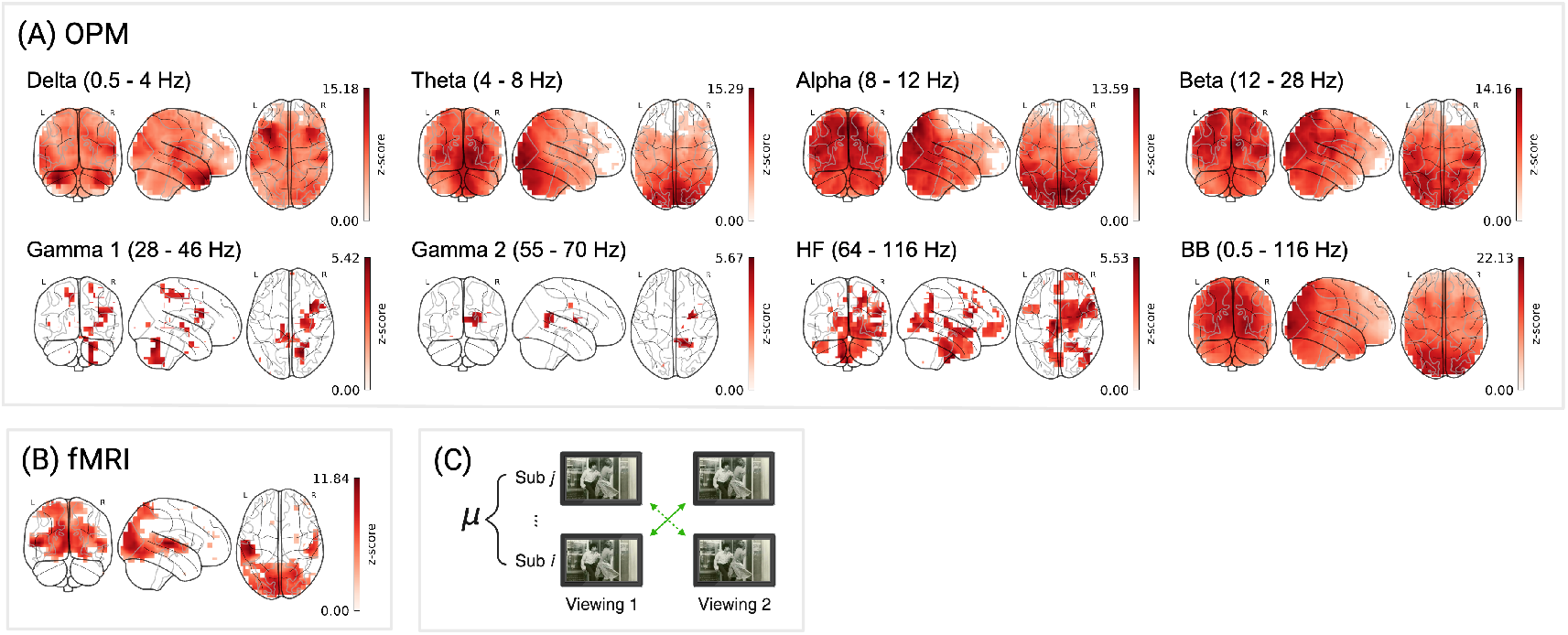
Between-subject reliability for OPM and fMRI. FDR-corrected *z*-scores are shown for: (A) OPM, 3,581 source locations across 10 subjects for 8 frequency bands; (B) fMRI: 3,581 voxels, across 11 subjects. (C) Schematic of between-subject correlation analysis. Color bars reflect *z*-scores (surviving correction for MC).

Interestingly, although reliability estimates were generally lower in the between-subject relative to the within-subject comparison for OPM, certain frequency bands exhibited the opposite pattern. Greater between-subject reliability can emerge because individual-specific noise contributes to within-subject testretest correlations. Idiosyncratic noise in between subject correlations may decrease to a larger extent than shared signal, effectively improving the SNR. For instance, regions reflecting a mixture of stimulus-driven and subject-specific processes may fail to reach significance within subjects yet emerge as significant between subjects once individual variance is attenuated. Most notably, we observed greater between-subject reliability in the *δ* and *β* bands, where peaks shifted from the cuneus to superior temporal and occipital areas, respectively. This is consistent with prior work demonstrating that ISCs during naturalistic movie viewing extend well beyond primary sensory regions, particularly to the superior temporal sulcus (Hasson et al., 2004). Moreover, these results align with previous findings of increased cross-subject synchronization in the *δ* band during emotionally salient scenes (Maffei, 2020).

For fMRI, although fewer voxels survived correction for MC (24.5%; Table 2, fMRI) compared to the within-subject analysis, the spatial pattern was largely preserved, with effects localized to auditory and visual regions (*M*_*z*_ = 4.51, *SD*_*z*_ = 1.97; see Figure 4B). The maximum was observed in the left superior temporal gyrus (peak *z* = 11.84), consistent with its role in processing complex audio-visual stimuli (Ghinst et al., 2016; Park et al., 2018; Ye et al., 2017).

### 3.3 Between-method comparisons

#### 3.3.1 OPM to fMRI

Consistent cross-modal alignment between OPM and fMRI was observed across all frequencies, with the most pronounced effects localized to visual processing regions in the mid-range and high-frequency bands (see Figure 5A). After correction for MC, peak *z*-scores ranged from -12.42 (*β*) to 5.19 (HF), with the largest proportion of surviving channels observed in the *α* band (17.8%; *M*_*z*_ = -4.37, *SD*_*z*_ = 3.05; Table 5, OPM to fMRI). Importantly, we observed negative cross-modal correlations in the *δ, θ, α*, and *β* bands, and positive correlations in the *γ*_1_, *γ*_2_, and high-frequency bands, in line with previous demonstrations of the relationship between MEG power spectra and BOLD signal (Zumer et al., 2010). Broadband reliability between fMRI and OPM closely resembled that of the lower frequencies, with peak effects observed in the superior occipital gyrus (peak *z*-score = -12.73; Table 5, OPM to fMRI).

**Figure 5:**
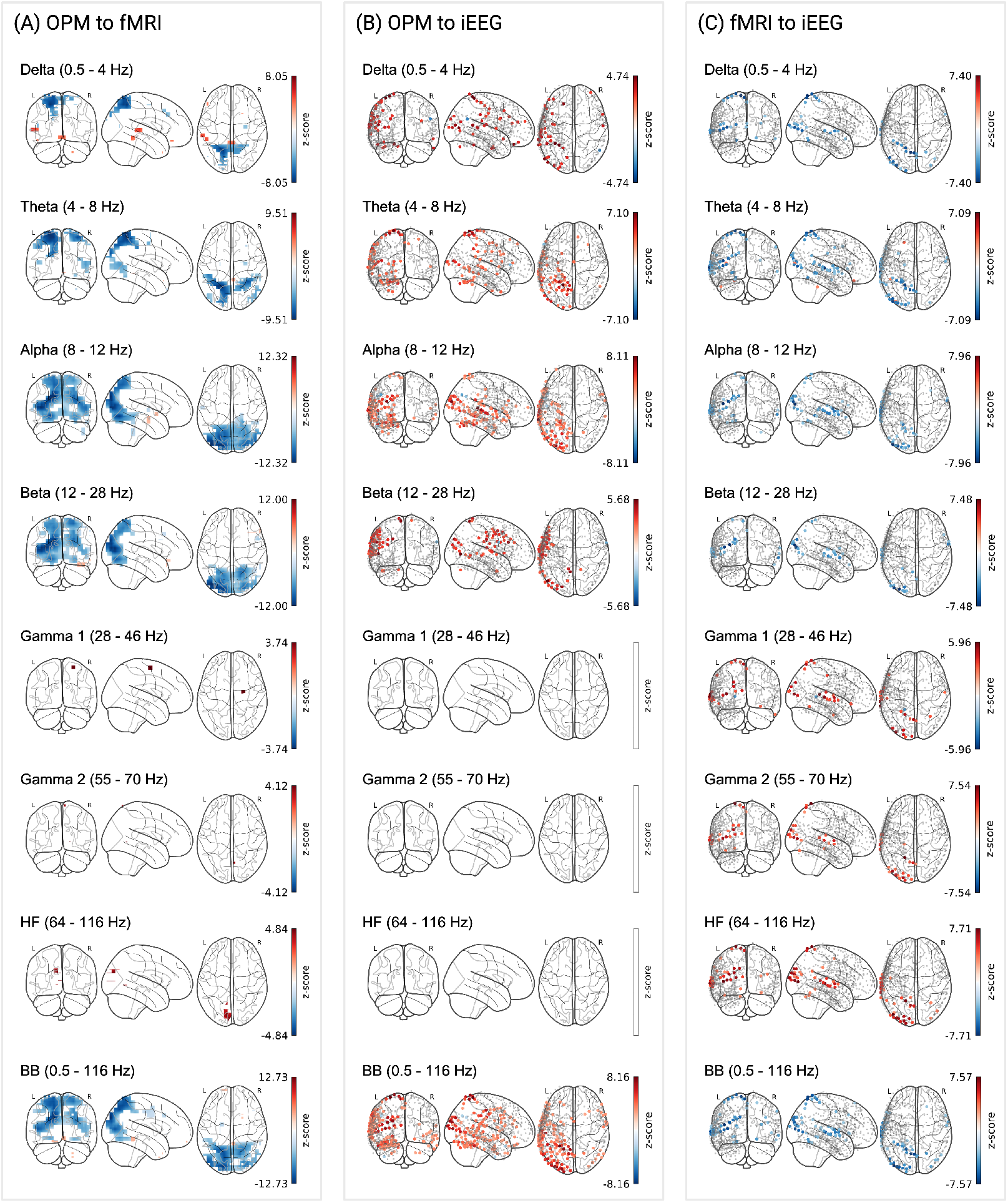
Consistent between-method alignment. (A) OPM to fMRI: *z*-scores from OPM subjects (n = 10) and fMRI subjects (n = 11) are shown for 3,581 locations. (B) OPM to iEEG: *z*-scores from OPM subjects (n = 10) and single-subject iEEG from 656 electrodes pooled across five subjects. (C) fMRI to iEEG: *z*-scores from fMRI subjects (n = 11) and single-subject iEEG from 656 electrodes pooled across five subjects. Color bars reflect *z*-scores (see above).

Of the frequency bands that demonstrated broader topographies (*δ, θ, α* and *β*), peak *z*-scores were located in the left superior parietal gyrus (*δ* and *θ*) and the left middle occipital gyrus (*α* and *β*), with a notable focal peak observed for the HF band in left cuneus. These spatial patterns are consistent with prior findings demonstrating that MEG and fMRI exhibit the greatest intersubject consistency in occipital areas during naturalistic movie viewing (Lankinen et al., 2018).

Though OPM-fMRI reliability patterns largely converged on regions reliable within each modality, supplemental analyses identified effects exclusive to each comparison (see Table S6 and S8). Interestingly, when contrasting within-subject OPM and within-subject fMRI, OPM-specific effects were observed in the cerebellum, providing convergent support for previous work demonstrating that OPM-MEG can capture cerebellar activity (Lin et al., 2019). This finding is notable given the prior claims that the architecture of the cerebellum restricts its contributions to SQUID-MEG signal, (Freeman et al., 2009) relative to fMRI; however, cerebellar effects in OPM may also reflect spatial leakage from visual regions due to limitations of the source reconstruction. Compared to SQUIDs, improved sensor-to-scalp proximity may afford OPMs greater sensitivity to cerebellar sources. Likewise, structures such as the basal ganglia have been shown to contribute weakly to SQUID-MEG signals due to their radial, non-columnar organization (Niedermeyer & da Silva, 2004). Yet, cross-modal-specific peaks emerged in regions of the basal ganglia for the OPM-fMRI comparison (Figure S7A). Additionally, prefrontal regions, which have previously exhibited weak intersubject coherence during naturalistic viewing (Hasson et al., 2004), demonstrated fMRI-specific reliability within subjects, but emerged as even more reliable in the OPM-fMRI comparison, with higher mean and maximum *z*-scores, as well as a larger proportion of channels surviving correction for MC (see Table S6). These patterns were largely consistent in the between-subject versus cross-modal contrasts, with one notable difference in the *δ* band, where OPM-specific peak reliability was observed in the temporal pole that was absent in between-subject fMRI. This finding may reflect continuous language comprehension during the movie that is uniquely captured with the greater temporal resolution of OPM (Mesulam, 2022). Though more spatially focal, these effects were also present in the *β* band of the OPM-fMRI comparison. Collectively, these results suggest that while each modality has distinct measurement strengths, cross-modal approaches can enhance sensitivity to shared activity that may be difficult to capture with either modality independently.

#### 3.3.2 OPM to iEEG

Cross-modal consistency between OPM and iEEG was more modest, with significant effects were observed in lower frequency bands, spanning frontal, parietal, and ventral visual regions (see Figure 5B). As reported in Table 5, OPM to iEEG, peak *z*-scores ranged from 4.74 (*δ*) to 8.11 (*α*), with the greatest proportion of correspondence observed in the *α* band (% MC = 16.3, *M*_*z*_ = 3.57, *SD*_*z*_ = 1.89; Table 5, OPM to iEEG).

Spatially, peak correspondence in the lower bands was localized to the superior parietal gyrus (*δ* and *θ*), middle temporal gyrus (*α*), and middle occipital gyrus (*β*). Although no effects were found in the *γ*_1_, *γ*_2_, and HF frequency bands, significant cross-modal consistency between OPM and iEEG was further observed in the broadband signal, with 24.2% of electrodes surviving correction for MC, and a consistent peak in the superior parietal gyrus (see Figure 5B).

When contrasting within-subject OPM and within-subject iEEG (see Table S7), OPM-specific reliability was broadly observed primarily in visual regions, whereas no regions were uniquely reliable in iEEG. The between-method OPM-iEEG comparison nevertheless revealed sparse, primarily frontal peaks (Figure S7B), mirroring the OPM-fMRI results and further supporting the idea that cross-modal analyses can reveal overlapping dynamics that are less apparent within individual modalities.

#### 3.3.3 fMRI to iEEG

To provide a comprehensive evaluation of the consistency between all three imaging modalities, we reproduced the results of Haufe et al. (2018) comparing iEEG to fMRI.

We observed significant cross-modal correlations between fMRI and iEEG across all frequency bands, with effects spanning temporal, parietal, and occipital regions (Figure 5C), consistent with Haufe et al. (2018). As in our fMRI to OPM comparison, negative correlations were observed in lower-frequency bands (*δ*-*β*), while (*γ*) and high-frequency bands showed positive associations with the BOLD signal (see Table 5, fMRI to iEEG). Peak *z*-scores ranged from -7.96 (*α*) to 7.71 (HF), with the largest proportion of channels surviving correction for MC in the *θ* band (7.6%, *M*_*z*_ = - 3.76, *SD*_*z*_ = 2.12).

Peak effects were largely located in areas associated with auditory and linguistic processing (Bhaya-Grossman & Chang, 2022; Wernicke, 1974), including the middle temporal(*θ* and *γ*_1_) and occipital (*α, β*, HF, and BB) gyri. This pattern is consistent with prior work demonstrating strong BOLD coupling to oscillatory activity in auditory cortex during movie viewing (Mukamel et al., 2005).

### 3.4 Signal-to-noise (SNR) comparisons

#### 3.4.1 Comparison of within-subject SNR to between-subject SNR in OPM

We observed clusters of significantly reduced between-subject SNR relative to within-subject SNR for OPM across all frequency bands (*p* < .001, controlling the false discovery rate at *q* < .025, as per cluster-based permutation; See methods). Maximum negative clusters were spatially distributed across sensorimotor, parietal, and frontal regions, reflecting areas with greater variability between subjects (Figure 6). The most pronounced SNR loss was observed in the *α* band, which yielded the largest significant negative cluster (85 channels, sum = -557.77), corresponding in the observed data to a region centered on the right postcentral gyrus (peak *z* = -8.84, *M*_*z*_ = -6.56, *SD*_*z*_ = 1.00). As shown in the top panel of Table 4, significant decreases in SNR were also found across the *δ* (sum = -313.80, max in inferior temporal gyrus), *θ* (sum = -72.94, max in cuneus), *β* (sum = -87.73, max in postcentral gyrus), *γ*_1_ (sum = -331.11, max in superior temporal gyrus), *γ*_2_ (sum = -283.24, max in gyrus rectus), HF (sum = -137.95, max in middle temporal gyrus), and broad bands (sum = -263.46, max in precentral gyrus).

**Table 4:**
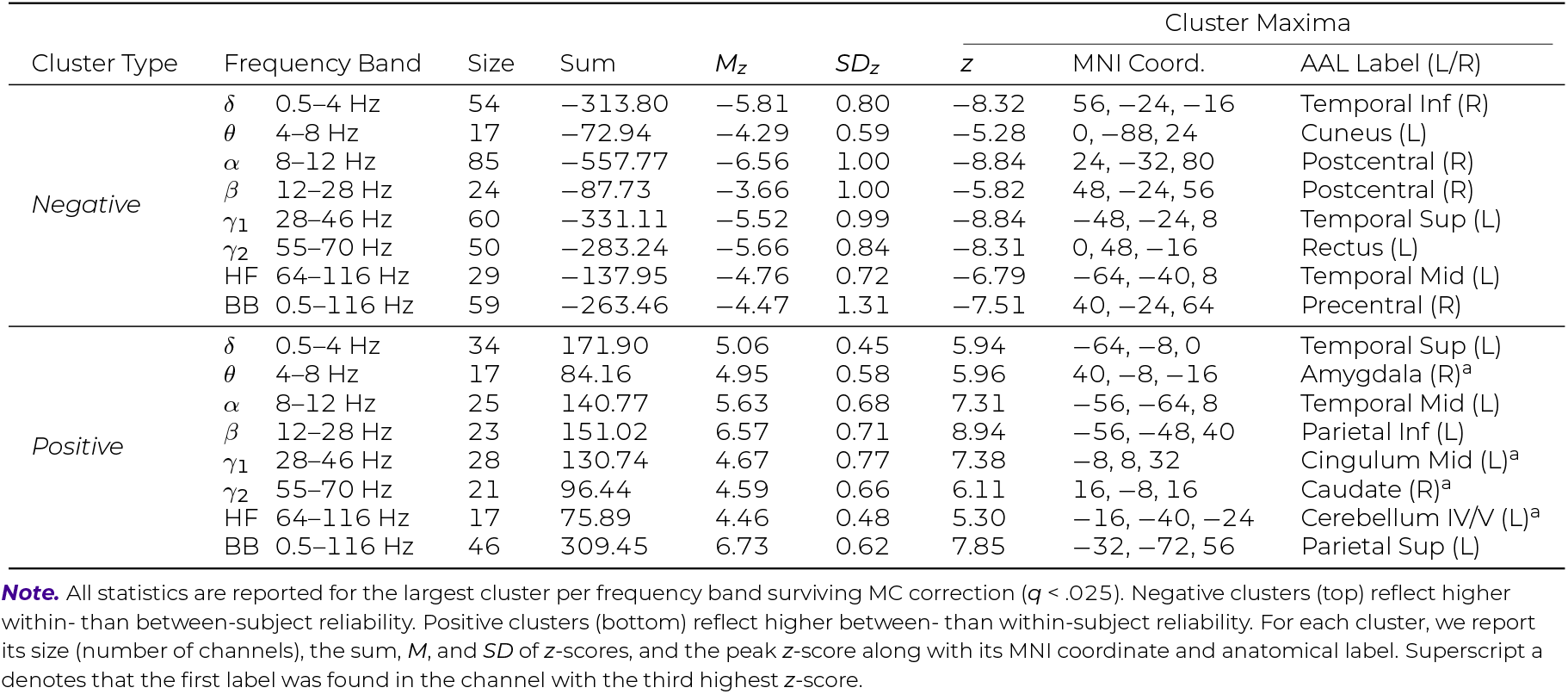
Cluster statistics for SNR differences in within- and between-subject OPM comparisons.

**Figure 6:**
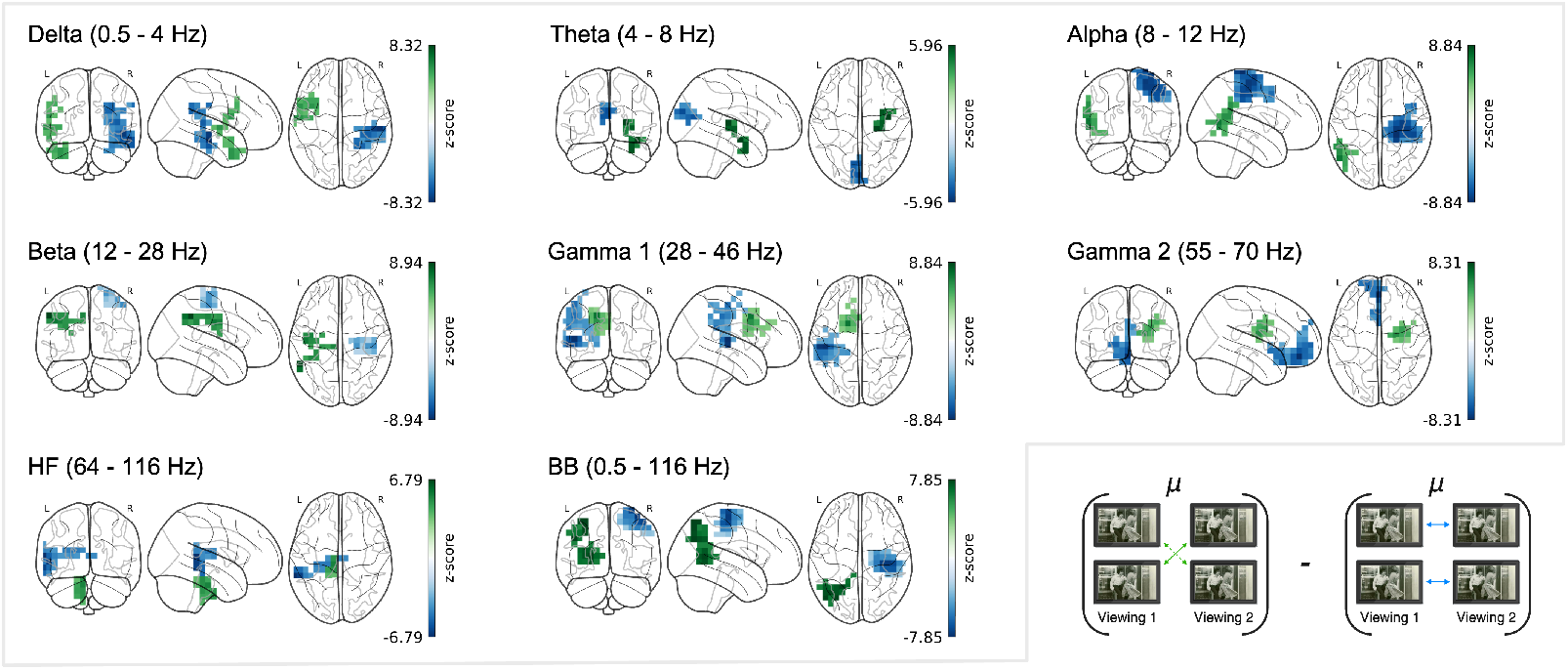
SNR differences in within- and between-subject comparisons for OPM. Within-modality SNR differences in OPM were computed by subtracting within-subject from between-subject *z*-scores at each channel. Negative values (blue) indicate greater signal consistency within individuals than across individuals; positive values (green) indicate greater signal consistency across individuals than within individuals. Significant clusters surviving MC correction (FDR *q* < .025) are outlined in the respective color. SNR changes are shown across frequency bands from *δ* (0.5- 4 Hz) to high-frequency (64-116 Hz), as well as the broadband (0.5–116 Hz). Diagrams below illustrate the comparison logic. Maps of uncorrected clusters identified by the initial cluster-forming threshold are shown in Figure S3A.

Additionally, all frequencies revealed clusters of significant positive SNR, representing areas where neural responses were more consistent across the OPM cohort than within individuals. These clusters were largely left-lateralized and predominately localized posterior association and limbic areas (see Figure 6), consistent with previous findings of intersubject synchrony during emotionally engaging movie viewing (Jääskeläinen et al., 2008; Maffei, 2020). Posterior peaks spanned the superior temporal (*δ*, sum = 171.90), middle temporal (*α*, sum = 140.77), inferior parietal (*β*, sum = 151.02), and superior parietal gyri (BB, sum = 309.45) (see Table 4). Additional positive clusters were found in subcortical regions, including *θ* (sum = 84.16, max in amygdala) and *γ*_2_ (sum = 96.44, max in caudate nucleus), as well as in the cerebellum (HF: sum = 75.89), and middle cingulate gyrus (*γ*_1_: sum = 130.74). Such higher SNR between different subjects can stem from a decrease in shared noise that exceeds the decrease in shared signal.

#### 3.4.2 SNR comparison of intersubject correlations in OPM and inter-method correlations between OPM and fMRI

Cluster-based permutation testing revealed significant negative clusters surviving MC correction in all frequency bands, indicating greater signal consistency within modality (OPM) than across modalities (OPM to fMRI). These effects were distributed primarily across frontal, parietal, temporal, and occipital regions (see Figure 7), with the largest cluster appearing in the *θ* band (84 channels, sum = -1000.63), corresponding to a region centered on the left lingual gyrus (*z* = -14.98, *M*_*z*_ = -11.91, *SD*_*z*_ = 0.94; Table 5, top panel). Across frequency bands, negative clusters mainly comprised posterior dorsal regions, including the precuneus (*α*, sum = -499.24), superior parietal gyrus (*β*, sum = - 568.91), and middle occipital gyrus (BB, sum = -1329.52), highlighting robust reliability within OPM that was not strongly reflected in fMRI.

**Figure 7:**
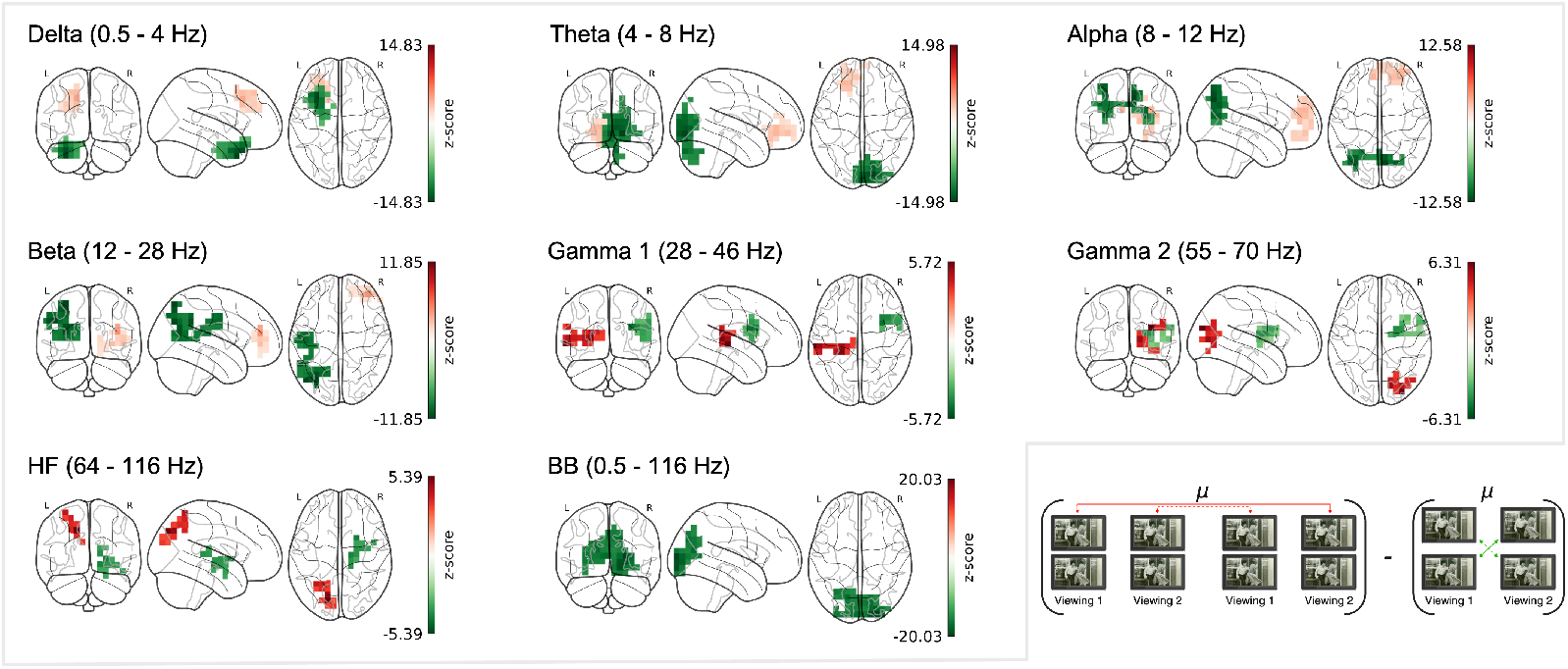
SNR differences within and across OPM and fMRI. Between-modality SNR differences contrast the SNR from between-subject OPM comparisons with the SNR from between-method OPM-to-fMRI comparisons. Negative values (green) indicate greater signal reliability within the OPM sample than in the cross-modal comparison. Positive values (red) indicate greater signal reliability in the cross-modal comparison (fMRI-to-OPM) than within the OPM sample. Significant clusters surviving MC correction (FDR *q* < .025) are outlined in the respective color. SNR changes are shown across frequency bands from *δ* (0.5-4 Hz) to high-frequency (64-116 Hz), as well as the broadband (0.5–116 Hz). Diagrams below illustrate the comparison logic for each panel. Maps of uncorrected clusters identified by the initial cluster-forming threshold are shown in Figure S3B.

**Table 5:** Cluster statistics for SNR differences in between-subject OPM and between-method OPM to fMRI comparisons.

| Cluster Type | Frequency Band | Size | Sum | $M_z$ | $SD_z$ | Cluster Maxima | | | |
| --- | --- | --- | --- | --- | --- | --- | --- | --- | --- |
| | | | | | | $z$ | MNI Coord. | AAL Label (L/R) | |
| Negative | $\delta$ 0.5–4 Hz | 43 | −456.72 | −10.62 | 1.69 | −14.83 | −32, 16, −40 | Temporal Pole Mid (L) | |
| | $\theta$ 4–8 Hz | 84 | −1000.63 | −11.91 | 0.94 | −14.98 | 16, −88, −8 | Lingual (R) | |
| | $\alpha$ 8–12 Hz | 49 | −499.24 | −10.19 | 0.95 | −12.58 | 8, −72, 40 | Precuneus (R) | |
| | $\beta$ 12–28 Hz | 57 | −568.91 | −9.98 | 0.70 | −11.85 | −32, −72, 56 | Parietal Sup (L) | |
| | $\gamma_1$ 28–46 Hz | 19 | −71.51 | −3.76 | 0.46 | −4.85 | 32, 16, 24 | Frontal Inf Tri (R)<sup>a</sup> | |
| | $\gamma_2$ 55–70 Hz | 26 | −88.11 | −3.39 | 0.60 | −4.70 | 64, 8, 24 | Precentral (R)<sup>a</sup> | |
|  | HF 64–116 Hz | 19 | −69.84 | −3.68 | 0.28 | −4.30 | 24, 8, 0 | Putamen (R) |  |
|  | BB 0.5–116 Hz | 83 | −1329.52 | −16.02 | 1.09 | −20.03 | 24, −96, 8 | Occipital Mid (R) |  |
| Positive | $\delta$ 0.5–4 Hz | 37 | 90.08 | 2.43 | 0.72 | 4.67 | −16, 24, 40 | Frontal Sup (L) | |
| | $\theta$ 4–8 Hz | 35 | 90.53 | 2.59 | 0.61 | 4.07 | −16, 56, −8 | Frontal Sup Orb (L)<sup>a</sup> | |
| | $\alpha$ 8–12 Hz | 37 | 91.56 | 2.47 | 0.45 | 3.45 | 32, 56, 24 | Frontal Mid (R) | |
| | $\beta$ 12–28 Hz | 24 | 46.09 | 1.92 | 0.86 | 4.57 | 40, 48, 16 | Frontal Mid (R) | |
| | $\gamma_1$ 28–46 Hz | 24 | 101.47 | 4.23 | 0.63 | 5.72 | −64, −24, 0 | Temporal Mid (L) | |
| | $\gamma_2$ 55–70 Hz | 18 | 81.95 | 4.55 | 0.60 | 6.31 | 32, −88, 24 | Occipital Mid (R) | |
|  | HF 64–116 Hz | 17 | 63.04 | 3.71 | 0.62 | 5.39 | −16, −80, 40 | Occipital Sup (L) |  |
|  | BB 0.5–116 Hz | — | — | — | — | — | — | — |  |

**Note.** All statistics are reported for the largest cluster per frequency band surviving MC correction ( $q < .025$ ). Negative clusters (top) reflect greater reliability between OPM subjects than when compared to fMRI. Positive clusters (bottom) reflect greater cross-modal correspondence between OPM and fMRI than within the OPM group alone. For each cluster, we report its size (number of channels), the sum, $M$ , and $SD$ of $z$ -scores, and the peak $z$ -score along with its MNI coordinate and anatomical label. Superscript a denotes that the first label was found in the channel with the second highest $z$ -score. Superscript b denotes that the first label was found in the channel with the third highest $z$ -score.

Surprisingly, significant positive clusters also emerged across all frequency bands, indicating that, in some regions, signal correspondence between OPM and fMRI exceeded within-modality reliability. In the lower frequencies, these clusters were largely concentrated in frontal regions (see Figure 7), likely indicating areas where activity is integrated over longer timescales, captured by both OPM and fMRI. Such clusters were observed in superior frontal (*δ* and *θ*) and middle frontal gyri (*α* and *β*). Moreover, and consistent with prior findings on high-frequency oscillations in visual processing (Martinovic & Busch, 2011; Mukamel et al., 2005), positive clusters were found in the occipital cortex in both the *γ*_2_ (sum = 81.95) and HF bands (sum = 63.04) (see Table 5, bottom panel), where, despite OPM’s fragmentary spatial pattern, alignment with fMRI was robust. Complementary to these effects, a positive cluster was identified in *γ*_1_, localized to the middle temporal gyrus (sum = 101.47), in line with proposed oscillatory dynamics during speech processing (Giraud & Poeppel, 2012). These findings indicate that, although within-modality reliability is generally higher, cross-modal correspondence may exceed it in anatomically and spectrally specific patterns.

## 4. DISCUSSION

We systematically evaluated the reliability of time-resolved neural signals in 8 frequency bands captured by OPM, and their cross-modal alignment with fMRI and iEEG during repeated viewings of a movie stimulus. OPM demonstrated widespread within-subject reliability, especially at lower frequencies; these results were consistent with SQUID-MEG studies of naturalistic movie viewing (Lankinen et al., 2018), but showed a broader spatial distribution that extended beyond occipital regions. We further observed reliable correlations *between* subjects in our OPM sample; their SNR exceeded, in some cases, that of within-subject analyses, which suggests that a shared signal consistently exceeded noise across individuals. Moreover, spatial maps became increasingly focused to sensory regions, mirroring patterns reported in previous intersubject analyses (Chang et al., 2015; Hasson et al., 2004; Haufe et al., 2018; Lankinen et al., 2014).

Correlations of OPM signals produced SNR estimates comparable in magnitude to those observed in fMRI and iEEG and with convergent spatial patterns in visual and auditory regions. OPM signals also directly correlated with both modalities, with the largest effects localized to visual areas. Cross-modal comparisons indicated that OPM showed slightly stronger alignment with fMRI than with iEEG in terms of peak *z*-score magnitude; however, mean *z*-scores and the proportion of channels surviving correction for MC were comparable, and sometimes greater between OPM and iEEG. Still, these differences may have been driven by differences in the number of channels in each comparison (i.e., OPM-fMRI: n = 3,581; OPM-iEEG: n = 656). Contrary to previous SQUID–MEG based findings (Lankinen et al., 2018), OPM achieved reliable correlations with fMRI, and showed frequency-dependent shifts in correlation-direction that are consistent with the inverse relationship between low-frequency oscillatory power and BOLD signals (Zumer et al., 2010). Collectively, these results demonstrate that OPM reliably captures stimulus-locked neural dynamics shared across individuals and imaging modalities.

Still, shared activity between two modalities does not imply uniform cross-modal congruence across all frequency bands. For instance, correspondence between all three modalities was comparable in the lower frequency range (LF; 0.5–28 Hz). At higher frequencies (HF; 28–116 Hz), however, only a few channels survived correction for multiple comparisons in the OPM-fMRI comparison, despite reliable iEEG-fMRI correspondence. This pattern may arise from frequency-dependent differences in the spatial and temporal sensitivity between OPM and iEEG. LF activity unfolds over longer timescales and spreads more broadly (Buzsáki & Draguhn, 2004), affording a higher SNR in the recorded signal. This high SNR in LFs likely supports the observed correspondence between OPM and iEEG, despite potential modality-specific sensitivities to sulcal and gyral sources.

In contrast, transient and local HF activity is reliably detected with intracranial recordings (Parvizi & Kastner, 2018). While direct comparisons between iEEG and SQUID-MEG are sparse and have largely been evaluated in clinical contexts, it has been shown that MEG reliably captures LF dynamics observed in iEEG, but nonetheless, has reduced sensitivity to highly localized highgamma activity and seizure onset zones (Dalal et al., 2009; Kim et al., 2016). This discrepancy may be driven by a combination of differential sensitivity to sulcal versus gyral sources, the limited spatial distribution of HF activity, and the proximity of intracranial electrodes to the neural sources that produce them. Thus, while OPM captures fragments of high-frequency activity, OPM and iEEG might not capture the same neural generators, which can be addressed in future studies. Likewise, the reduced magnitude of HF OPM-fMRI correspondence may be at least partially attributed to the lower within-modality reliability in OPM compared to other frequency bands.

Although reliability within individual modalities was generally convergent between them, supplemental analyses revealed modality-specific effects as well as regions that emerged as reliable only in cross-modal comparisons. Across analyses, OPM exhibited modality-specific sensitivity to regions that are typically difficult to detect with SQUID-MEG, including the cerebellum, consistent with prior work (Lin et al., 2019) and likely reflecting improved sensor proximity and temporal resolution. In contrast, within-subject OPM–iEEG comparisons showed OPM-exclusive reliability primarily in visual regions, with no regions uniquely reliable in iEEG, consistent with the idea that OPM captures large-scale synchronized activity that may not dominate individual intracranial contacts. Several regions, such as prefrontal cortex and deep structures, were weak or inconsistent in within-modality contrasts but emerged more reliably in OPM-fMRI and OPM-iEEG comparisons, suggesting that cross-modal alignment enhances sensitivity to shared activity.

We note, however, that spatial leakage inherent to MEG source reconstruction warrants caution in interpreting the spatial extent of these effects. Consistent with this, although reliability patterns were largely consistent across source reconstruction approaches, supplemental analyses using minimum-norm-estimation yielded less focal spatial distributions. As such, we recommend viewing these reliability patterns as heatmaps of where neural sources are commensurable with the measured data and focusing on peak locations as the most plausible generators of the observed signals. More broadly, this underscores that within-modality reliability alone is insufficient to establish that the measured signal represents meaningful neural activity, highlighting the importance of cross-modal comparisons. Our results demonstrate that OPM signal is not only reliable within and across individuals, but critically, that it captures neural activity measured with independent imaging modalities.

One advantage of our analysis approach is that it directly quantifies the reliability of shared signals relative to shared noise. In our analyses, Pearson correlation coefficients (*r*-values) increased substantially when down-sampling to 0.67 Hz. The computed *z*-scores, on the other hand, account for temporal autocorrelation and noise structure, offering an appropriate metric of signal reliability relative to surrogate data (Haufe et al., 2018). Furthermore, by conducting pairwise analyses at a common sampling rate of 20 Hz, we preserved intersubject variance and temporal precision and were able to demonstrate the reliability of OPM both within and between individuals. Significant effects emerged despite minimal preprocessing and early-generation sensor hardware, which underscores the efficacy of OPM that can likely be improved with future advances in sensor design and denoising techniques (Rier et al., 2023).

Importantly, estimating SNR via *z*-scores places all modalities on a shared statistical scale, allowing for comparisons across datasets with differing temporal resolution, spatial coverage, and sample sizes. Leveraging this approach, we observed that within-subject analyses yielded more widespread reliability than between-subject comparisons. Indeed, between-subject comparisons appeared spatially focused in primary sensory regions, consistent with neural responses commonly evoked during movie viewing (Hasson et al., 2004; Lankinen et al., 2014). I.e., while it is often considered a limitation of a design when data are acquired from separate participant groups, because of the introduction of subject-specific variability into comparisons, our findings suggest that intersubject analyses can also mitigate idiosyncratic biases by attenuating shared noise (e.g., spatial leakage in source reconstruction) and task-unrelated fluctuations. In our data, between-subject comparisons effectively isolated stimulus-driven responses generalizable across individuals (Hasson et al., 2004, 2010; Nastase et al., 2019). We acknowledge that residual misalignment arising from spatial normalization may reduce anatomical correspondence, and that cross-modal reliability with iEEG may be further affected by partial volume effects in fMRI at cortical boundaries (Ballester et al., 2002) or depth bias in OPM source reconstruction. However, because such effects are expected to attenuate rather than inflate correlation estimates, the reported results should be interpreted as a lower bound on shared signal. Our SNR comparisons further suggest that this principle can extend across imaging modalities. Speculatively, within-subject designs with cross-modal comparisons may also overestimate reliability by capturing stable but characteristic individual signal components such as subject-specific lateralization.

Finally, we observed that SNR can – in some regions – increase when comparisons are drawn between different subjects undergoing different imaging modalities. While differences in modality sensitivity often reduce overall alignment, they may also support the suppression of shared noise and can consequently sometimes lead to increased SNR. In particular, while high-frequency OPM activity appeared spatially diffuse, its correlation with fMRI was tightly localized to regions expected to support visual processing. I.e., while high-frequency measurements in OPM may not be stable enough to be reliably localized across subjects or measurements within OPM, its similarity to fMRI signal allowed for the demonstration that it is reliably present. Notably, such preserved shared signal can only be detected in regions where the modalities functionally align. In the future, cross-modal convergence may be leveraged to guide denoising and source reconstruction, effectively integrating the spatial precision of fMRI with the temporal resolution of MEG.

In conclusion, our findings demonstrate that during audio-visual processing, OPM achieves reliability levels comparable to fMRI and iEEG; these effects were most robust in low-frequency bands. OPM showed strong correspondence with BOLD signals, following the established patterns of inverse relationships between high- and low-frequency neural activity. These results underscore the potential of OPM as a flexible, noninvasive tool capable of resolving complex brain activity across multiple temporal and spatial scales.

## DATA AND CODE AVAILABILITY

Upon publication of this manuscript, all data and analysis code will be made openly available at https://osf.io/urjvb.

## AUTHOR CONTRIBUTIONS

Olivia R. Christiano: Conceptualization; Data curation; Formal analysis; Project administration; Software; Visualization; Writing – original draft. Sebastian Michelmann: Conceptualization; Data curation; Formal analysis; Methodology; Resources; Software; Supervision; Writing – review & editing.

## DECLARATION OF COMPETING INTERESTS

The authors declare that there are no competing interests.

## ACKNOWLEDGMENTS

We would like to thank Uri Hasson and Christopher J. Honey, who generously shared the data that made this work possible, as well as Stefan Haufe, who made the preprocessed fMRI data from Arcaro et al. (2015) available and pioneered the analysis approach. We also thank Lukas Rier and coauthors, notably Elena Boto and Matthew J. Brookes, for making the OPM data publicly available and for valuable comments on early conceptualization and analyses of this study.

## SUPPLEMENTARY MATERIAL

Supplementary material for this article is included at the end of this document.

## Supplementary Material

**Table S1:** Comparison of within-subject maximum correlations after temporally down-sampling and averaging.

| Modality | Frequency | Maxima | | | $r$ (20 Hz) | $r$ (0.67 Hz) | $r$ (0.67 Hz, GA) |
| --- | --- | --- | --- | --- | --- | --- | --- |
| | | $z$ | MNI Coord. | AAL Label (Left/Right) | | | |
| OPM | $\delta$ 0.5–4 Hz | 13.42 | 8, –96, 16 | Cuneus (R) | .08 | .19 | .61 |
| | $\theta$ 4–8 Hz | 16.04 | 8, –96, 16 | Cuneus (R) | .08 | .20 | .55 |
| | $\alpha$ 8–12 Hz | 13.57 | 0, –88, 32 | Cuneus (L) | .08 | .23 | .63 |
| | $\beta$ 12–28 Hz | 12.28 | –8, –96, 24 | Cuneus (L) | .04 | .22 | .65 |
| | $\gamma_1$ 28–46 Hz | 6.01 | –40, –16, 64 | Precentral (L) | .02 | .12 | .42 |
| | $\gamma_2$ 55–70 Hz | 5.85 | 0, 56, 0 | Frontal Sup Medial (L) | .02 | .12 | .30 |
|  | HF 64–116 Hz | 5.28 | 40, 56, 8 | Frontal Mid (R) | .02 | .11 | .43 |
|  | BB 0.5–116 Hz | 20.76 | 0, –96, 24 | Cuneus (L) | .07 | .30 | .66 |
| fMRI | – – | 13.75 | –56, –16, 0 | Temporal Mid (L) | .35 | .32 | .71 |
| iEEG | $\delta$ 0.5–4 Hz | 12.35 | –58, –14, 48 | Postcentral (L) | .29 | .58 | – |
| | $\theta$ 4–8 Hz | 9.79 | 65, –18, 25 | SupraMarginal (R) | .22 | .66 | – |
| | $\alpha$ 8–12 Hz | 10.75 | –68, –11, 8 | Temporal Sup (L) | .24 | .59 | – |
| | $\beta$ 12–28 Hz | 9.74 | –67, –21, 16 | SupraMarginal (L) | .11 | .74 | – |
| | $\gamma_1$ 28–46 Hz | 7.80 | –48, –54, –24 | Temporal Inf (L) | .09 | .73 | – |
| | $\gamma_2$ 55–70 Hz | 8.30 | 65, –20, 15 | Temporal Sup (R) | .12 | .51 | – |
|  | HF 64–116 Hz | 21.26 | –67, –21, 16 | SupraMarginal (L) | .34 | .86 | – |
|  | BB 0.5–116 Hz | 12.30 | –68, –11, 8 | Temporal Sup (L) | .25 | .82 | – |

**Note.** For each imaging modality and frequency band, we identify the maximum within-subject $z$ -score (surviving multiple comparisons correction) and report correlation coefficients at that location. First, we report the untransformed correlation at the peak $z$ -score location using data sampled at 20 Hz ( $r$ (20 Hz)), where correlations were computed between the first and second movie viewing for each subject and channel and averaged across subjects. Next, we report the corresponding value from the same location using data that were further down-sampled to 0.67 Hz ( $r$ (0.67 Hz)). Finally, for OPM and fMRI, we averaged the 0.67 Hz time series across subjects separately for viewing 1 and viewing 2, computed the correlation between the two grand-averaged signals, and report the resulting untransformed coefficient ( $r$ (0.67, GA)). For fMRI, analyses were conducted on data projected to the OPM source grid for consistency. $r$ (0.67, GA) is not applicable to the single-subject iEEG data.

### Within-viewing correlation analyses

For within-viewing comparisons, we followed the same procedures described in the main Methods section, except we correlated time series from the same viewing (e.g., subject *i*’s first viewing with subject *j*’s first viewing) rather than between them. Correlations were computed separately for viewing 1 and viewing 2, then averaged prior to generating surrogate distributions and computing *z*-scores.

**Table S2:**
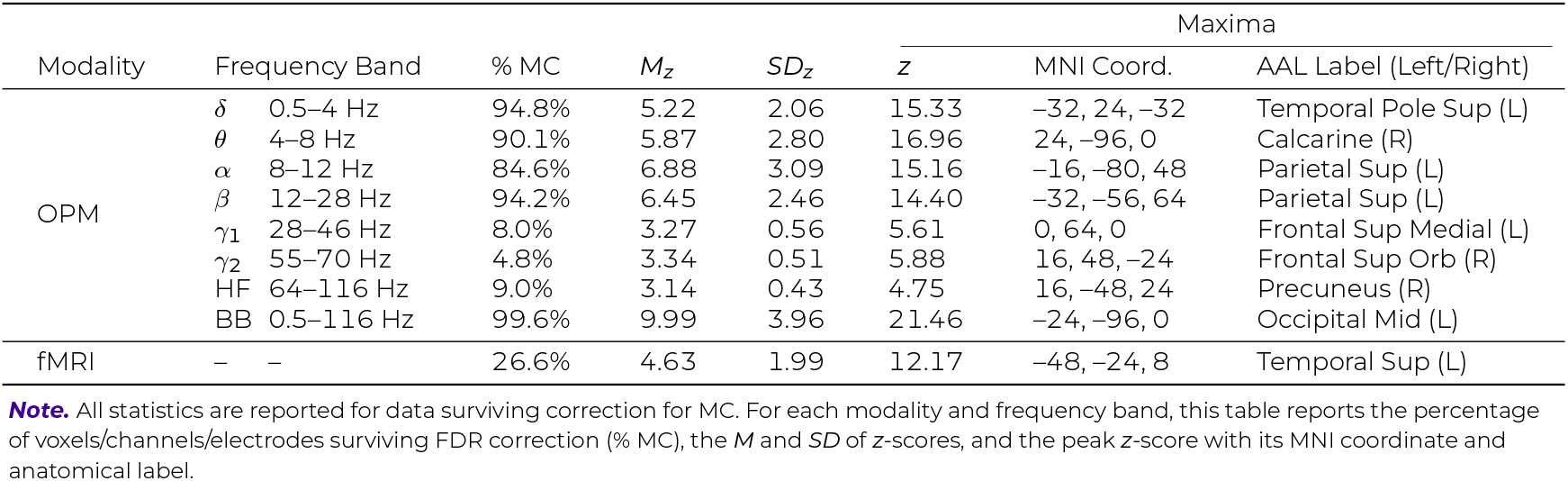
Between-subject reliability statistics for OPM and fMRI within movie viewings.

**Figure S1:**
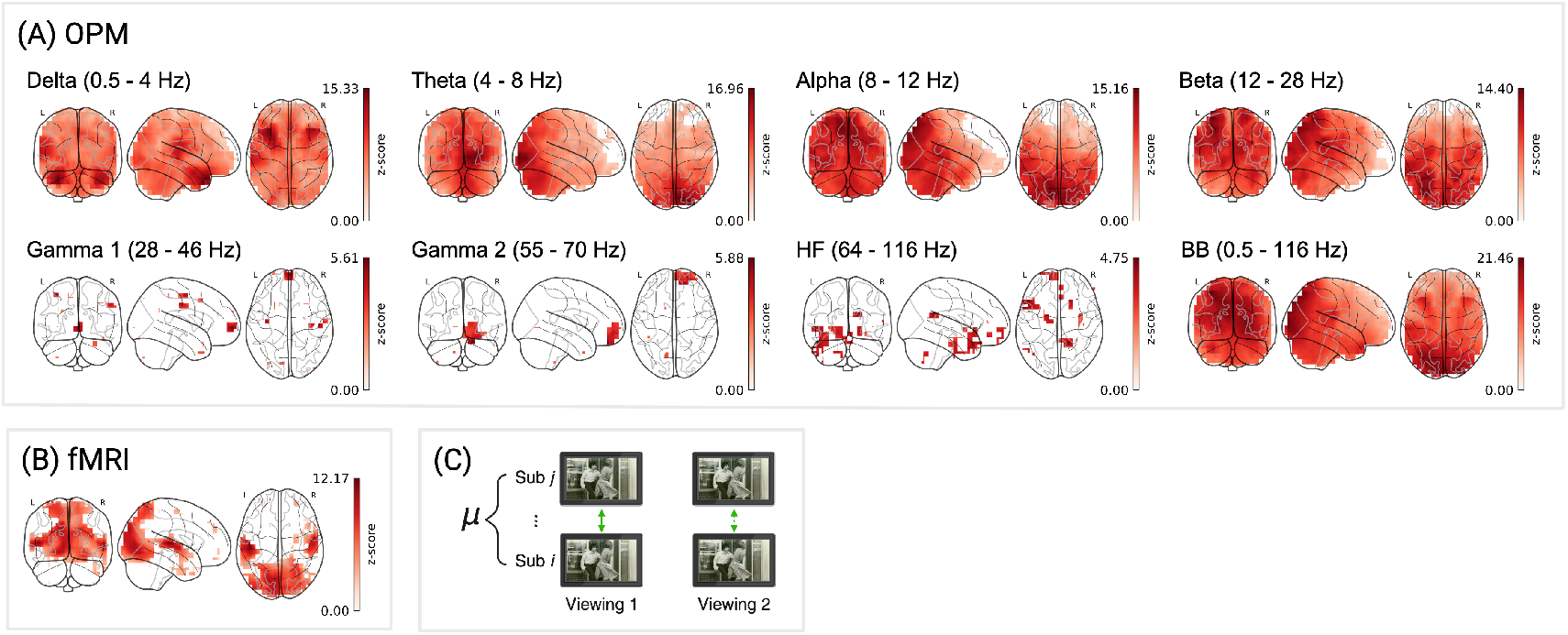
Between-subject reliability for OPM and fMRI within movie viewings. FDR-corrected *z*-scores are shown for: (A) OPM, 3,581 source locations across 10 subjects for 8 frequency bands; (B) fMRI: 3,581 voxels, across 11 subjects. (C) Schematic of between-subject correlation analysis. Color bars reflect *z*-scores (surviving correction for MC).

**Table S3:** Between-method reliability statistics across frequency bands for OPM, fMRI, and iEEG within movie viewings.

| Comparison | Frequency Band | % MC | $M_z$ | $SD_z$ | Maxima | | |
| --- | --- | --- | --- | --- | --- | --- | --- |
| | | | | | $z$ | MNI Coord. | AAL Label (Left/Right) |
| OPM to fMRI | $\delta$ 0.5–4 Hz | 4.8% | −0.83 | 3.83 | −7.42 | −32, −56, 56 | Parietal Sup (L) |
| | $\theta$ 4–8 Hz | 11.7% | −2.25 | 3.71 | −11.46 | −24, −56, 72 | Parietal Sup (L) |
| | $\alpha$ 8–12 Hz | 16.2% | −4.40 | 3.12 | −13.26 | −40, −88, 8 | Occipital Mid (L) |
| | $\beta$ 12–28 Hz | 16.1% | −4.22 | 2.91 | −12.03 | −24, −88, 24 | Occipital Sup (L) |
| | $\gamma_1$ 28–46 Hz | 0.0% | 3.85 | 0.00 | 3.85 | −24, −8, 64 | Frontal Sup (L) |
| | $\gamma_2$ 55–70 Hz | 1.0% | 3.17 | 2.58 | 5.74 | 24, −88, 0 | Calcarine (R) |
|  | HF 64–116 Hz | 1.1% | 3.41 | 2.64 | 6.61 | 0, −96, 24 | Cuneus (L) |
|  | BB 0.5–116 Hz | 13.7% | −3.88 | 3.55 | −11.69 | −16, −56, 64 | Precuneus (L) |
| OPM to iEEG | $\delta$ 0.5–4 Hz | 3.5% | 3.41 | 1.58 | 5.45 | −21, −55, 78 | Parietal Sup (L) |
| | $\theta$ 4–8 Hz | 7.3% | 3.13 | 2.35 | 7.29 | −21, −55, 78 | Parietal Sup (L) |
| | $\alpha$ 8–12 Hz | 12.2% | 3.83 | 1.58 | 8.00 | −35, −91, 22 | Occipital Mid (L) |
| | $\beta$ 12–28 Hz | 8.1% | 3.63 | 1.28 | 6.91 | −24, −87, 33 | Occipital Sup (L) |
| | $\gamma_1$ 28–46 Hz | 0.0% | – | – | – | – | – |
| | $\gamma_2$ 55–70 Hz | 0.0% | – | – | – | – | – |
|  | HF 64–116 Hz | 0.0% | – | – | – | – | – |
|  | BB 0.5–116 Hz | 24.2% | 3.67 | 1.66 | 8.41 | −21, −55, 78 | Parietal Sup (L) |
| fMRI to iEEG | $\delta$ 0.5–4 Hz | 4.6% | −3.44 | 2.02 | −5.43 | −46, −83, 7 | Occipital Mid (L) |
| | $\theta$ 4–8 Hz | 8.1% | −3.83 | 1.74 | −7.16 | −74, −28, 3 | Temporal Mid (L) |
| | $\alpha$ 8–12 Hz | 5.3% | −4.74 | 1.27 | −7.41 | −67, −8, 3 | Temporal Sup (L) |
| | $\beta$ 12–28 Hz | 5.3% | −4.01 | 0.90 | −6.73 | −24, −87, 33 | Occipital Sup (L) |
| | $\gamma_1$ 28–46 Hz | 2.3% | 4.28 | 2.35 | 6.67 | −39, −90, 10 | Occipital Mid (L) |
| | $\gamma_2$ 55–70 Hz | 4.0% | 3.97 | 2.56 | 7.26 | −39, −90, 10 | Occipital Mid (L) |
|  | HF 64–116 Hz | 6.7% | 4.50 | 1.88 | 8.08 | −24, −87, 33 | Occipital Sup (L) |
|  | BB 0.5–116 Hz | 8.4% | −4.22 | 1.07 | −6.75 | −74, −28, 3 | Temporal Mid (L) |

**Note.** All statistics are reported for data surviving correction for MC. For each comparison and band, we report the percent of voxels/electrodes surviving correction for MC (% MC), the $M$ and $SD$ of $z$ -scores, and the peak $z$ -score with its MNI coordinate and anatomical label.

**Figure S2:**
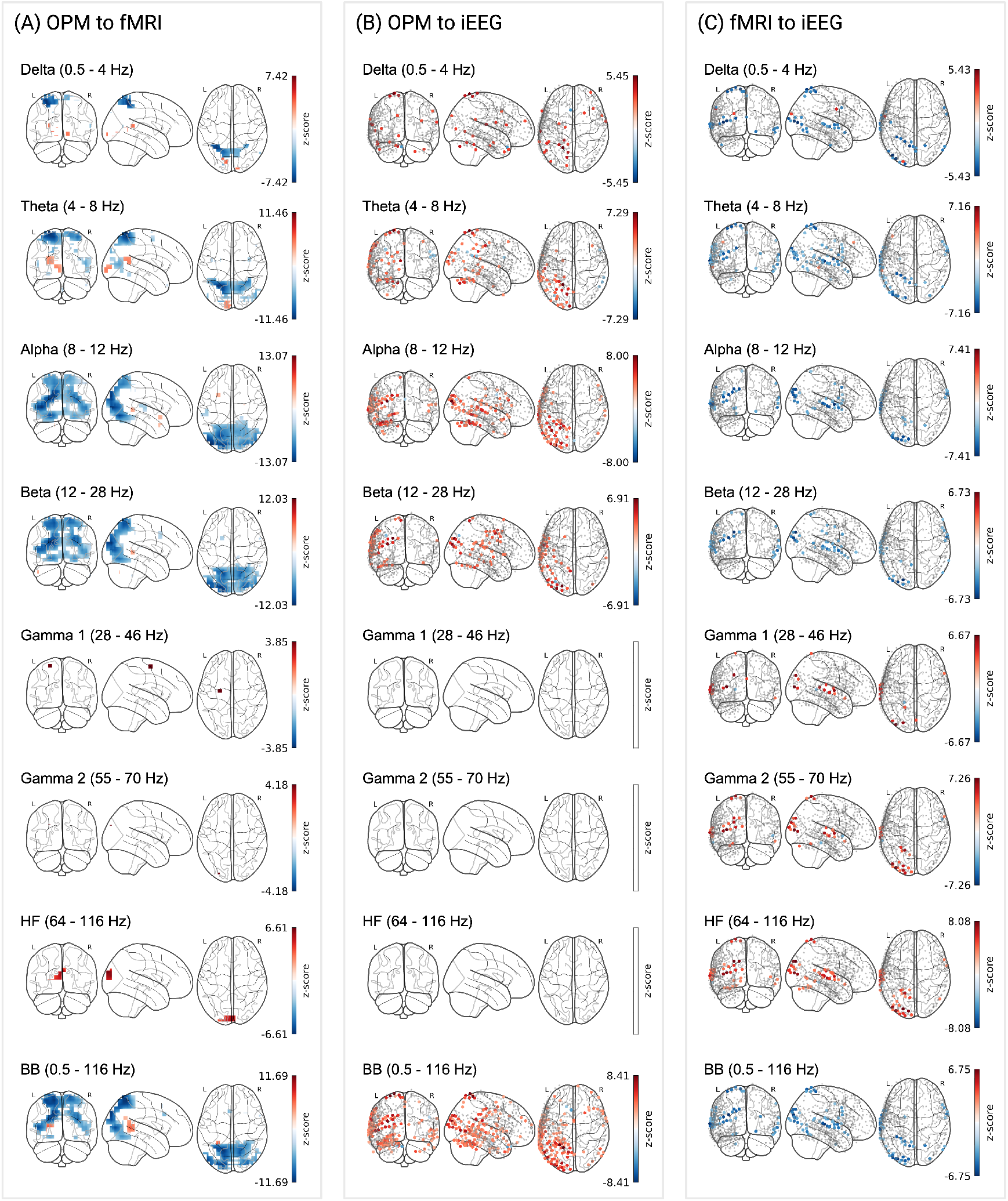
Consistent between-method alignment within movie viewings. (A) OPM to fMRI: *z*-scores from OPM subjects (n = 10) and fMRI subjects (n = 11) are shown for 3,581 locations. (B) OPM to iEEG: *z*-scores from OPM subjects (n = 10) and single-subject iEEG from 656 electrodes pooled across five subjects. (C) fMRI to iEEG: *z*-scores from fMRI subjects (n = 11) and single-subject iEEG from 656 electrodes pooled across five subjects. Color bars reflect *z*-scores (see above).

**Figure S3:**
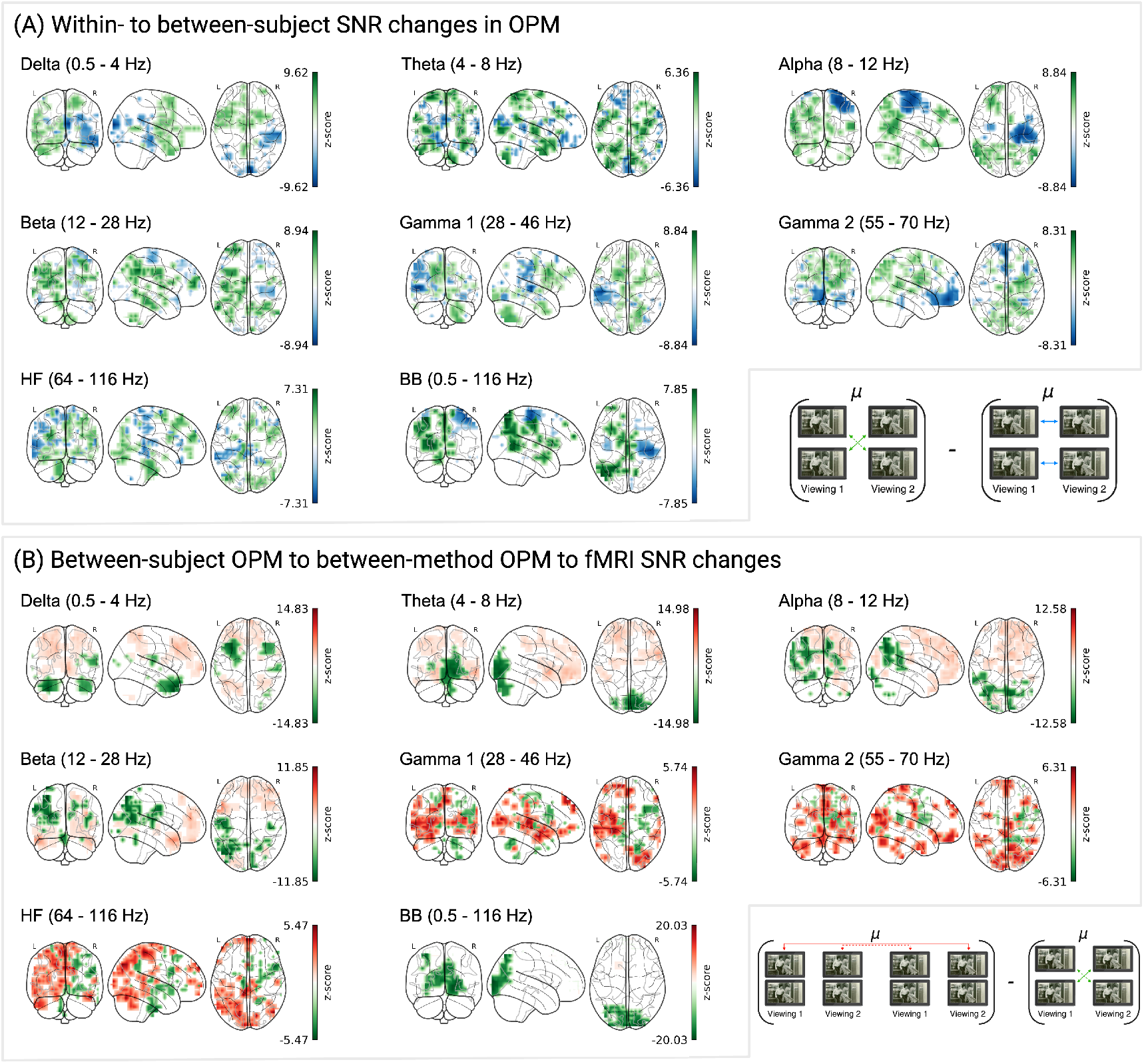
Uncorrected cluster maps of SNR differences within and across modalities. All clusters shown exceed the 97.5th or fall below the 2.5th percentile of the empirical SNR distribution; values were not corrected for multiple comparisons and do not reflect statistical significance. (A) Within-modality SNR changes in OPM, computed by subtracting within-subject from between-subject *z*-scores at each channel. Negative values (blue) indicate greater signal consistency within individuals than across individuals. (B) Between-modality SNR changes comparing between-subject OPM to between-method fMRI, where negative values (red) indicate greater signal reliability within OPM than in cross-modal correspondence. SNR changes are shown across frequency bands from delta (0.5- 4 Hz) to high-frequency (64-116 Hz). Diagrams below illustrate the comparison logic for each panel.

### Supplemental analyses with alternate source reconstruction parameters

To ensure that the reliability patterns we observed for OPM were not dependent on the specific source reconstruction parameters used in our main analyses, we repeated all OPM analyses using an alternative inverse solution. Specifically, while our primary results were obtained using LCMV beamforming, we additionally employed a minimum-norm-estimation (MNE) approach, which does not consider the data covariance.

Across source reconstruction approaches, OPM exhibited broadly consistent reliability patterns. Using LCMV beamforming (Table 1, OPM; Table 2, OPM), within- and between-subject reliability was most pronounced in the broadband and lower frequency bands, with effects concentrated in visual and temporal processing regions, while higher-frequency bands showed sparser and more spatially restricted patterns. In our supplemental analyses using an MNE solution (Table S4) reliability estimates were slightly reduced in magnitude overall, and demonstrated less focal spatial distributions, consistent with the smoother nature of minimum-norm estimates (see Figure S4). Nevertheless, similar patterns were observed across source reconstruction methods, indicating that the observed reliability patterns are robust to source reconstruction choices.

**Table S4:** Within and between-subject reliability statistics for OPM with alternate source reconstruction parameters.

| Modality | Frequency Band | % MC | $M_z$ | $SD_z$ | Maxima | | | AAL Label (Left/Right) |
| --- | --- | --- | --- | --- | --- | --- | --- | --- |
| | | | | | $z$ | MNI Coord. | | |
| within-subject | $\delta$ 0.5–4 Hz | 67.7% | 3.62 | 1.32 | 10.84 | 48, –16, 40 | | Postcentral (R) |
| | $\theta$ 4–8 Hz | 84.1% | 3.57 | 1.19 | 8.11 | 8, –96, 0 | | Calcarine (R) |
| | $\alpha$ 8–12 Hz | 91.4% | 5.24 | 1.97 | 9.80 | 0, 8, 40 | | Cingulum Mid (L) |
| | $\beta$ 12–28 Hz | 81.8% | 3.83 | 1.35 | 8.25 | –56, –24, 0 | | Temporal Mid (L) |
| | $\gamma_1$ 28–46 Hz | 30.8% | 3.02 | 0.70 | 5.66 | –48, –8, 32 | | Postcentral (L) |
| | $\gamma_2$ 55–70 Hz | 28.7% | 2.88 | 0.55 | 5.31 | 64, –40, –24 | | Temporal Inf (R) |
|  | HF 64–116 Hz | 63.1% | 2.99 | 0.71 | 6.76 | –56, –48, 48 |  | Parietal Inf (L) |
|  | BB 0.5–116 Hz | 99.1% | 4.78 | 1.57 | 10.11 | 48, –40, 56 |  | Parietal Inf (R) |
| between-subject | $\delta$ 0.5–4 Hz | 54.9% | 4.00 | 1.45 | 8.76 | –64, –16, 32 | | Postcentral (L) |
| | $\theta$ 4–8 Hz | 88.8% | 4.03 | 1.39 | 9.73 | 16, –88, –32 | | Crus II (R) |
| | $\alpha$ 8–12 Hz | 60.5% | 3.75 | 1.34 | 8.36 | –8, –72, 64 | | Precuneus (L) |
| | $\beta$ 12–28 Hz | 99.8% | 5.21 | 1.42 | 9.75 | –40, –64, –16 | | Fusiform (L) |
| | $\gamma_1$ 28–46 Hz | 15.6% | 2.99 | 0.45 | 4.50 | 56, –16, 48 | | Postcentral (R) |
| | $\gamma_2$ 55–70 Hz | 15.3% | 3.20 | 0.61 | 5.40 | 56, –48, –8 | | Temporal Mid (R) |
|  | HF 64–116 Hz | 45.4% | 3.01 | 0.62 | 5.50 | 16, 8, 8 |  | Caudate (R) |
|  | BB 0.5–116 Hz | 97.5% | 5.36 | 1.71 | 12.65 | –8, –72, 64 |  | Precuneus (L) |

**Note.** All statistics are reported for data surviving correction for MC. For each modality and frequency band, this table reports the percentage of voxels/channels/electrodes surviving FDR correction (% MC), the $M$ and $SD$ of $z$ -scores, and the peak $z$ -score with its MNI coordinate and anatomical label.

**Figure S4:**
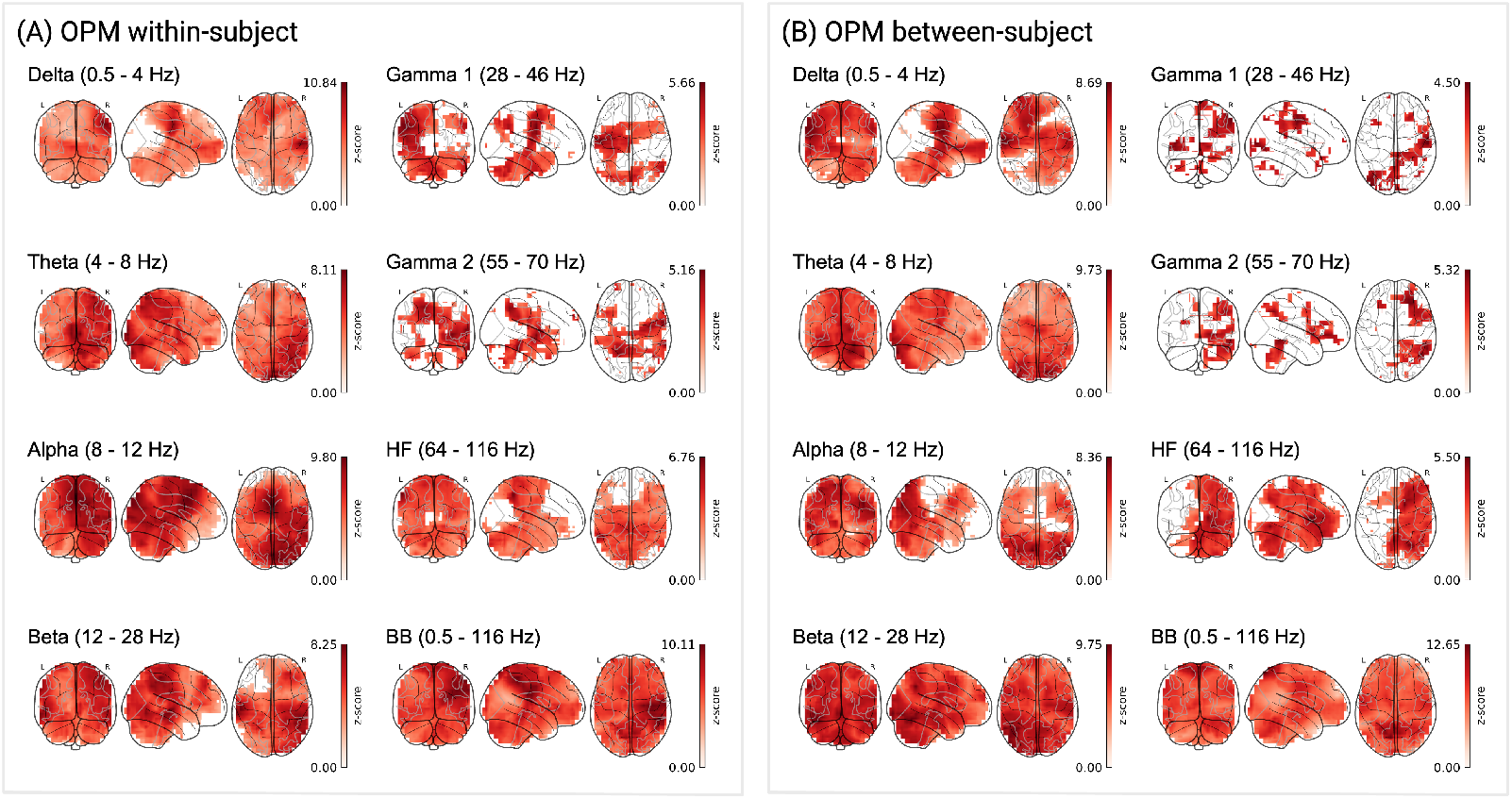
Within and between-subject reliability for OPM with alternate source reconstruction parameters. FDR-corrected *z*-scores are shown for OPM, computed at 3,581 source locations across 10 subjects for 8 frequency bands: (A) within-subject and (B) between-subject comparisons between the first and second movie viewing. Color bars reflect *z*-scores (surviving correction for MC).

A similar pattern was observed for the between-method comparisons (Table 3). Cross-modal reliability between OPM and fMRI, and between OPM and iEEG, using the alternate, MNE reconstruction (Table S5), moderately reduced *z*-score magnitudes, but sometimes increased their spatial extent (see Figure S5). Still, the observed patterns were maintained across both inverse solutions. Together, these findings indicate that the observed patterns of reliability in OPM reflect robust, stimulus-locked structure rather than artifacts of source reconstruction choices, though spatial interpretability may be reduced.

**Table S5:** Between-method reliability statistics for OPM, fMRI, and iEEG with alternate source reconstruction parameters.

| Comparison | Frequency Band | % MC | $M_z$ | $SD_z$ | $z$ | Maxima | |
| --- | --- | --- | --- | --- | --- | --- | --- |
|  |  |  |  |  |  | MNI Coord. | AAL Label (Left/Right) |
| OPM to fMRI | $\delta$ 0.5–4 Hz | 0.6% | −3.88 | 1.68 | −5.01 | 24, −8, 64 | Frontal Sup (R) |
| | $\theta$ 4–8 Hz | 11.4% | −2.80 | 3.32 | −8.23 | −16, −64, 64 | Parietal Sup (L) |
| | $\alpha$ 8–12 Hz | 9.3% | −3.79 | 2.19 | −8.95 | 48, −80, 0 | Occipital Mid (R) |
| | $\beta$ 12–28 Hz | 12.1% | −3.40 | 2.95 | −9.31 | −40, −88, 8 | Occipital Mid (L) |
| | $\gamma_1$ 28–46 Hz | — | — | — | — | — | — |
| | $\gamma_2$ 55–70 Hz | 2.1% | 3.78 | 0.97 | 5.51 | 16, −88, 24 | Cuneus (R) |
|  | HF 64–116 Hz | 3.6% | 4.00 | 1.19 | 6.69 | −8, −80, 32 | Cuneus (L) |
|  | BB 0.5–116 Hz | 8.5% | −3.46 | 2.28 | −7.10 | −16, −64, 64 | Parietal Sup (L) |
| OPM to iEEG | $\delta$ 0.5–4 Hz | 1.7% | 3.47 | 0.21 | 3.91 | 30, 61, −16 | Frontal Mid Orb (R) |
| | $\theta$ 4–8 Hz | 7.5% | 3.43 | 1.57 | 5.66 | −29, −49, 73 | Parietal Sup (L) |
| | $\alpha$ 8–12 Hz | 3.4% | 3.08 | 2.19 | 4.88 | −35, −91, 22 | Occipital Mid (L) |
| | $\beta$ 12–28 Hz | 4.4% | 3.59 | 1.46 | 5.44 | −8, −56, 70 | Precuneus (L) |
| | $\gamma_1$ 28–46 Hz | 0.2% | 4.11 | 0.00 | 4.11 | −65, 15, 5 | Frontal Inf Oper (L) |
| | $\gamma_2$ 55–70 Hz | — | — | — | — | — | — |
|  | HF 64–116 Hz | — | — | — | — | — | — |
|  | BB 0.5–116 Hz | 17.1% | 3.42 | 0.95 | 5.84 | −58, −22, 47 | Parietal Inf (L) |

**Note.** All statistics are reported for data surviving correction for MC. For each comparison and band, we report the percent of voxels/electrodes surviving correction for MC (% MC), $M$ and $SD$ of $z$ -scores, and the peak $z$ -score with its MNI coordinate and anatomical label.

**Figure S5:**
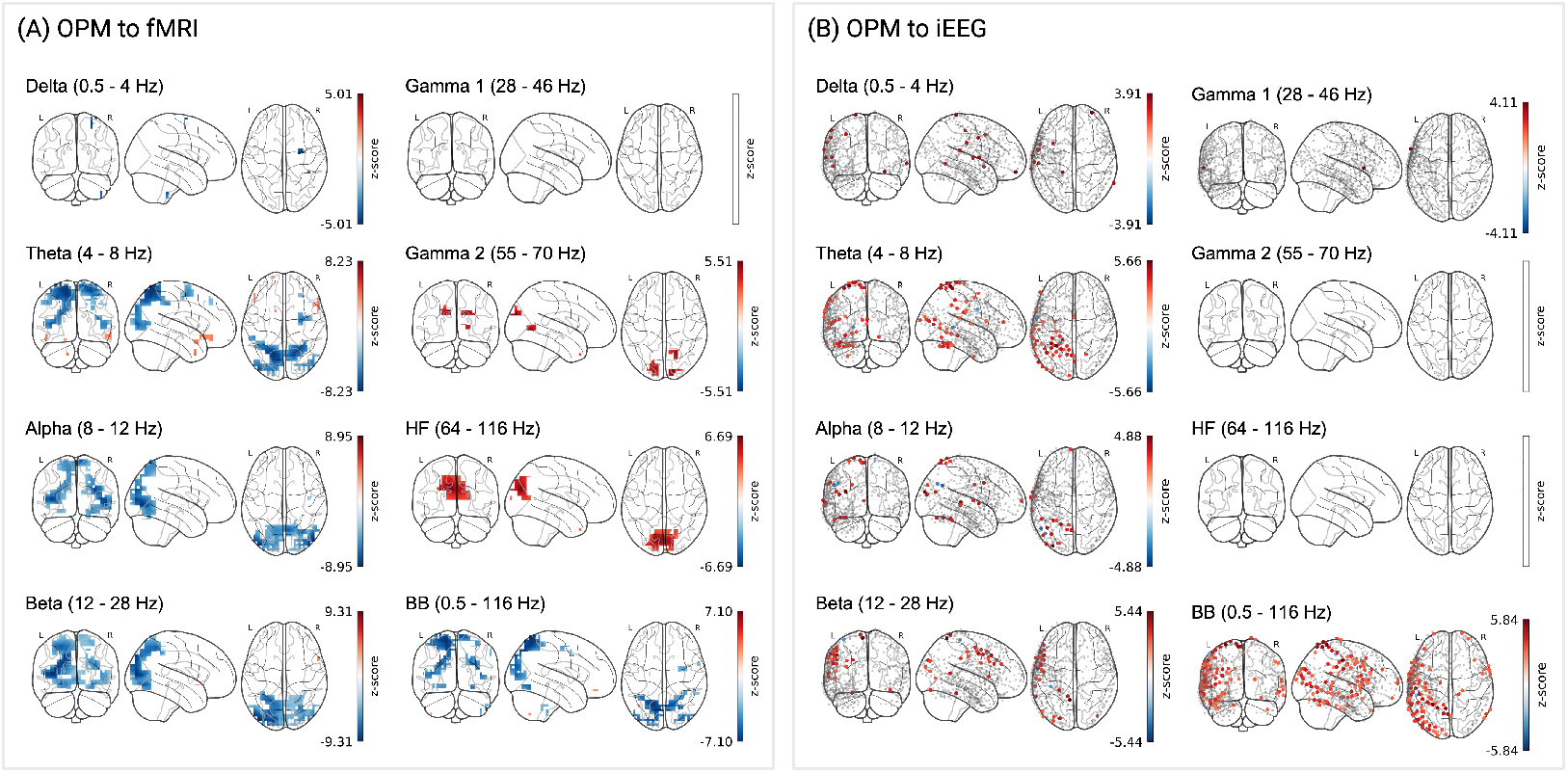
Consistent between-method alignment with alternate source reconstruction parameters. (A) OPM to fMRI: *z*-scores from OPM subjects (n = 10) and fMRI subjects (n = 11) are shown for 3,581 locations. (B) OPM to iEEG: *z*-scores from OPM subjects (n = 10) and single-subject iEEG from 656 electrodes pooled across five subjects. Color bars reflect *z*-scores (see above).

### Modality-specific vs cross-modal specific patterns of reliability

In addition to quantifying shared signal across modalities, we identified patterns of reliability exclusive to individual modalities, and regions that emerged as reliable only in cross-modal comparisons. Modality-specific patterns index the distinct sensitivities of each method, including potential differences in depth sensitivity, spatial specificity, and temporal precision. In contrast, cross-modal-specific patterns may reveal stimulus-driven signal that becomes detectable only when modality-specific noise is attenuated (i.e., there is less shared noise between modalities than within them). Examining these patterns therefore clarifies both what is uniquely captured by each modality and what may be missed when relying on any single methodological approach.

To address this, we performed overlap analyses of cross-modal comparisons for OPM to fMRI, and OPM to iEEG, that separated channels into three mutually exclusive categories: (i) channels showing reliable signal within one modality but not the other (e.g., significant *z*-scores in OPM that were not significant in fMRI), (ii) channels showing reliable signal within the second modality but not the first (e.g., significant *z*-scores in fMRI that were not significant in OPM), and (iii) channels that emerged as reliable only in between-method comparisons (e.g., OPM-fMRI) despite not surviving correction within either modality alone. To allow for direct comparison of iEEG within-subject reliability to OPM, we recomputed within-subject analyses using the OPM data that was reconstructed at the 656 iEEG electrode locations from cross-modal analyses. Binary reliability masks of *z*-scores that survived MC correction were created across each comparison and modality, combined to isolate modality-specific and cross-modal-specific channels, and projected into MNI space for visualization. We compared within-subject reliability to between-method reliability across modality pairs, and repeated the same analysis for the between-subject OPM-fMRI comparison.

**Table S6:**
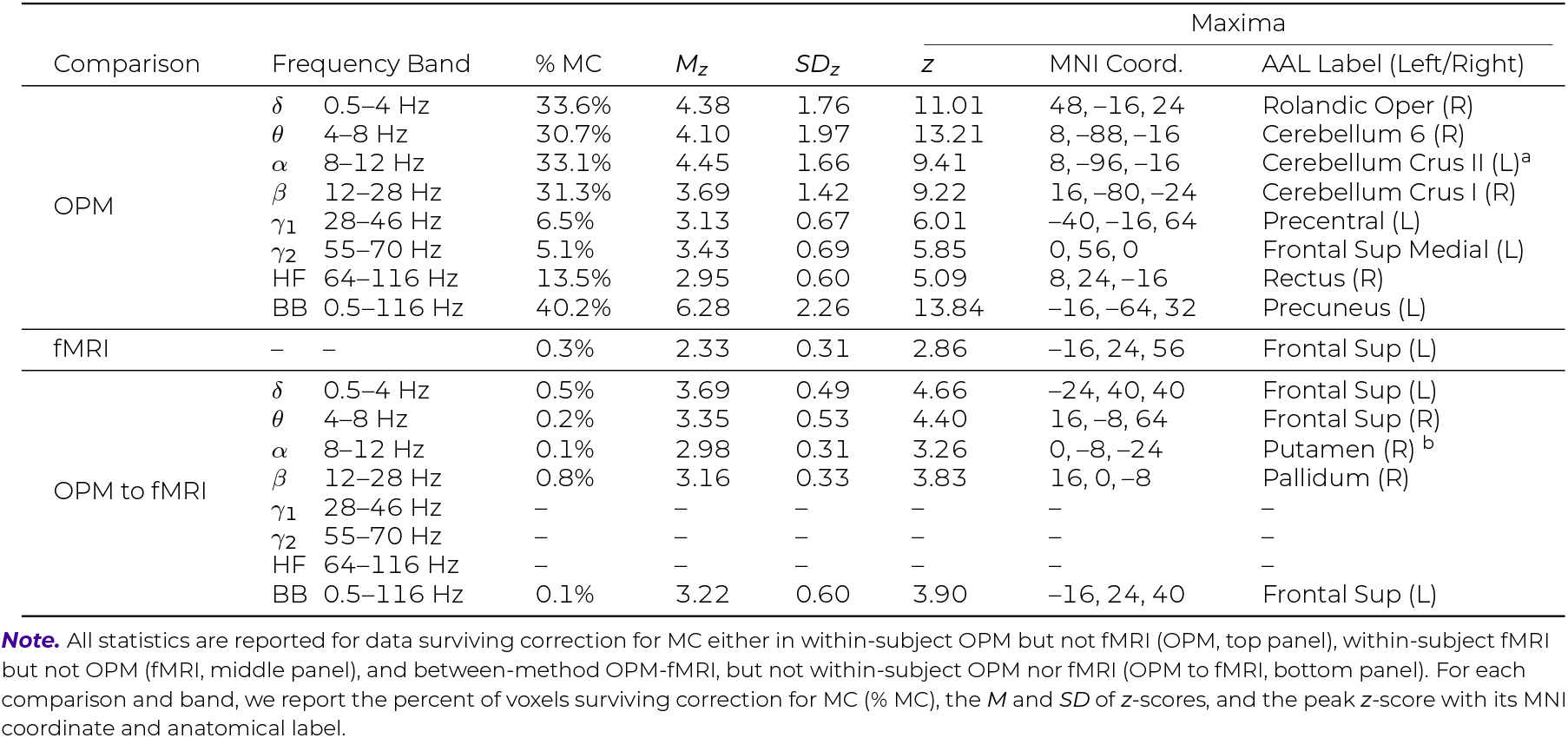
Patterns of reliability exclusive to within-subject OPM, within-subject fMRI, and between-method OPM-fMRI.

**Table S7:** Patterns of reliability exclusive to within-subject OPM, within-subject iEEG, and between-method OPM–iEEG.

| Comparison | Frequency Band | % MC | $M_z$ | $SD_z$ | Maxima | | | AAL Label (Left/Right) |
| --- | --- | --- | --- | --- | --- | --- | --- | --- |
|  |  |  |  |  | z | MNI Coord. |  |  |
| OPM | $\delta$ 0.5–4 Hz | 50.8% | 4.88 | 2.04 | 11.21 | -7, -99, 16 | | Cuneus (L) |
| | $\theta$ 4–8 Hz | 40.4% | 3.83 | 2.02 | 13.66 | 2, -81, 26 | | Cuneus (L) |
| | $\alpha$ 8–12 Hz | 61.9% | 4.76 | 1.86 | 12.01 | -7, -99, 16 | | Cuneus (L) |
| | $\beta$ 12–28 Hz | 56.7% | 4.07 | 1.73 | 9.88 | -21, -101, 15 | | Occipital Sup (L) |
| | $\gamma_1$ 28–46 Hz | 11.0% | 3.31 | 0.68 | 5.23 | 27, -24, 79 | | Precentral (R) |
| | $\gamma_2$ 55–70 Hz | 3.4% | 3.67 | 0.53 | 4.74 | 56, 9, -3 | | Temporal Pole Sup (R) |
|  | HF 64–116 Hz | 8.5% | 3.10 | 0.55 | 4.67 | 51, 32, 13 |  | Frontal Inf Tri (R) |
|  | BB 0.5–116 Hz | 53.2% | 6.37 | 2.83 | 19.89 | -3, -91, 32 |  | Cuneus (L) |
| iEEG | $\delta$ 0.5–4 Hz | – | – | – | – | – | | – |
| | $\theta$ 4–8 Hz | – | – | – | – | – | | – |
| | $\alpha$ 8–12 Hz | – | – | – | – | – | | – |
| | $\beta$ 12–28 Hz | – | – | – | – | – | | – |
| | $\gamma_1$ 28–46 Hz | – | – | – | – | – | | – |
| | $\gamma_2$ 55–70 Hz | – | – | – | – | – | | – |
|  | HF 64–116 Hz | – | – | – | – | – |  | – |
|  | BB 0.5–116 Hz | – | – | – | – | – |  | – |
| OPM to iEEG | $\delta$ 0.5–4 Hz | 0.6% | 3.44 | 0.46 | 4.12 | -61, 9, 29 | | Precentral (L) |
| | $\theta$ 4–8 Hz | 0.9% | 3.22 | 0.26 | 3.58 | 1, -8, 55 | | Supp Motor Area (R) |
| | $\alpha$ 8–12 Hz | 0.3% | 2.96 | 0.05 | 3.00 | -35, 65, 10 | | Frontal Sup (L) |
| | $\beta$ 12–28 Hz | 0.9% | 3.16 | 0.29 | 3.52 | -65, -55, 1 | | Temporal Mid (L) |
| | $\gamma_1$ 28–46 Hz | – | – | – | – | – | | – |
| | $\gamma_2$ 55–70 Hz | – | – | – | – | – | | – |
|  | HF 64–116 Hz | – | – | – | – | – |  | – |
|  | BB 0.5–116 Hz | – | – | – | – | – |  | – |

**Figure S6:**
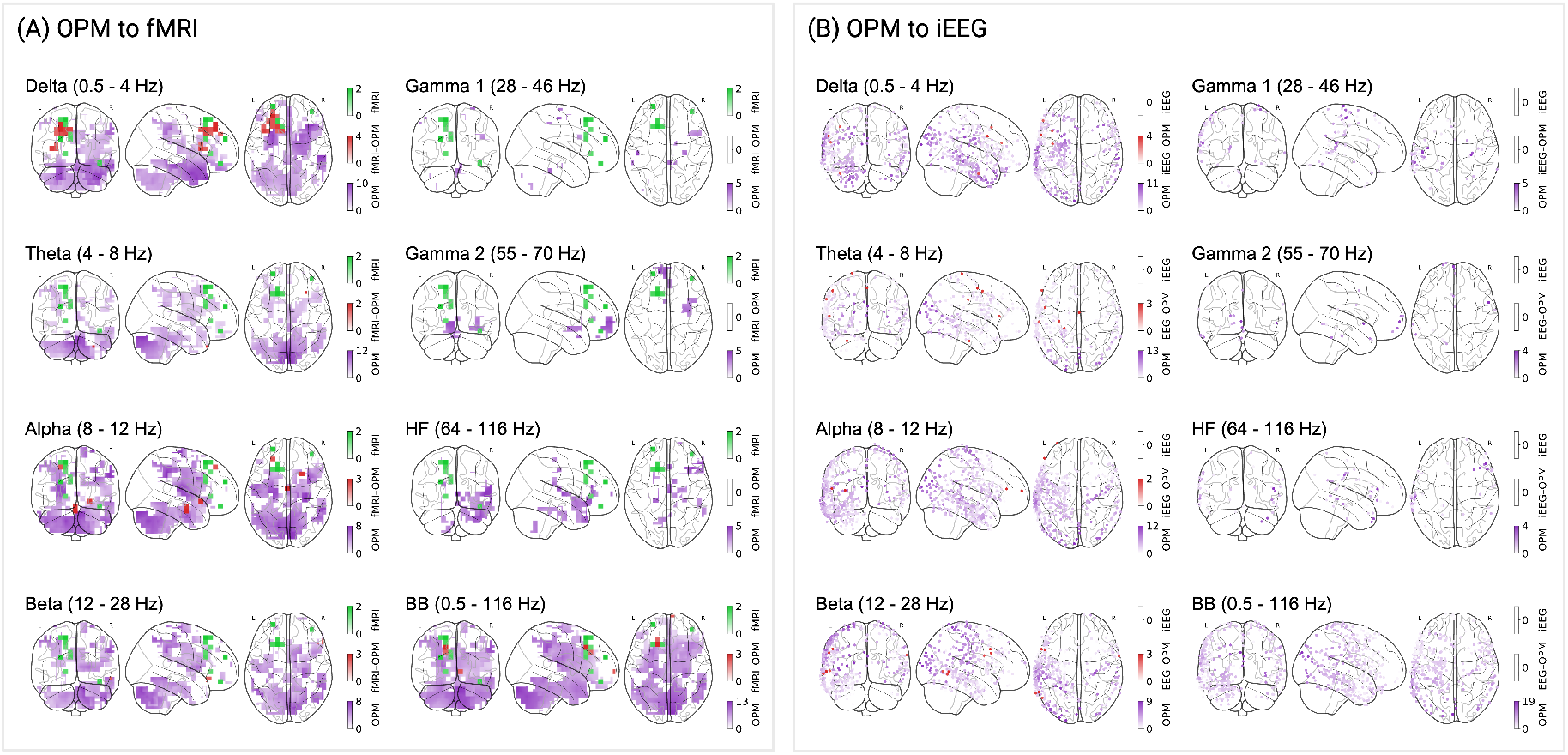
Reliable *z*-scores exclusive to within-subject and between-method comparisons across OPM, fMRI, and iEEG. Channels/electrodes surviving MC correction for (A) OPM to fMRI: within-subject OPM but not within-subject fMRI (purple), within-subject fMRI but not within-subject OPM (green), and between-method OPM-fMRI but not within-subject OPM nor fMRI (red), and (B) OPM to iEEG: within-subject OPM but not within-subject iEEG (purple), within-subject iEEG but not in within-subject OPM (note, no electrodes survived correction for MC within iEEG only for any frequency), and between-method OPM-iEEG but not in within-subject OPM nor iEEG (red). Color bars reflect *z*-scores (see above).

**Table S8:** Patterns of reliability exclusive to between-subject OPM, between-subject fMRI, and between-method OPM-fMRI.

| Comparison | Frequency Band | % MC | $M_z$ | $SD_z$ | Maxima | | | |
| --- | --- | --- | --- | --- | --- | --- | --- | --- |
| | | | | | $z$ | MNI Coord. | AAL Label (Left/Right) | |
| OPM | $\delta$ 0.5–4 Hz | 64.5% | 4.50 | 1.99 | 15.18 | –32, 24, –32 | Temporal Pole Sup (L) | |
| | $\theta$ 4–8 Hz | 59.4% | 4.74 | 2.32 | 13.50 | 8, –80, –40 | Cerebellum Crus II (R) | |
| | $\alpha$ 8–12 Hz | 56.5% | 5.37 | 2.32 | 13.59 | 8, –72, 56 | Precuneus (R) | |
| | $\beta$ 12–28 Hz | 69.4% | 5.61 | 2.16 | 12.52 | –32, –80, 40 | Occipital Mid (L) | |
| | $\gamma_1$ 28–46 Hz | 9.6% | 3.34 | 0.65 | 5.55 | 16, –64, –32 | Cerebellum 6 (R) | |
| | $\gamma_2$ 55–70 Hz | 3.5% | 3.52 | 0.62 | 5.67 | 16, –48, 16 | Precuneus (R) | |
|  | HF 64–116 Hz | 20.2% | 2.98 | 0.62 | 5.50 | 56, 16, 24 | Frontal Inf Tri (R) |  |
|  | BB 0.5–116 Hz | 75.0% | 8.14 | 2.99 | 18.20 | –32, –72, 56 | Parietal Sup (L) |  |
| fMRI | – | – | – | – | – | – | – |  |
| OPM to fMRI | $\delta$ 0.5–4 Hz | 0.3% | 3.52 | 0.51 | 4.59 | –24, 40, 48 | Frontal Sup (L) | |
| | $\theta$ 4–8 Hz | 0.2% | 3.25 | 0.60 | 4.43 | –32, –8, 56 | Precentral (L) | |
| | $\alpha$ 8–12 Hz | 0.1% | 2.92 | 0.17 | 3.16 | 24, –32, 48 | Postcentral (R) | |
| | $\beta$ 12–28 Hz | 0.4% | 3.38 | 0.54 | 4.23 | –48, 16, –24 | Temporal Pole Sup (L) | |
| | $\gamma_1$ 28–46 Hz | – | – | – | – | – | – | |
| | $\gamma_2$ 55–70 Hz | – | – | – | – | – | – | |
|  | HF 64–116 Hz | 0.0% | 3.48 | 0.00 | 3.48 | –48, –8, 16 | Postcentral (L) |  |
|  | BB 0.5–116 Hz | – | – | – | – | – | – |  |

**Note.** All statistics are reported for data surviving correction for MC either in between-subject OPM but not fMRI (OPM, top panel), between-subject fMRI but not OPM (fMRI, middle panel), and between-method OPM-fMRI, but not between-subject OPM nor fMRI (OPM to fMRI, bottom panel). For each comparison and band, we report the percent of voxels surviving correction for MC (% MC), the $M$ and $SD$ of $z$ -scores, and the peak $z$ -score with its MNI coordinate and anatomical label.

**Figure S7:**
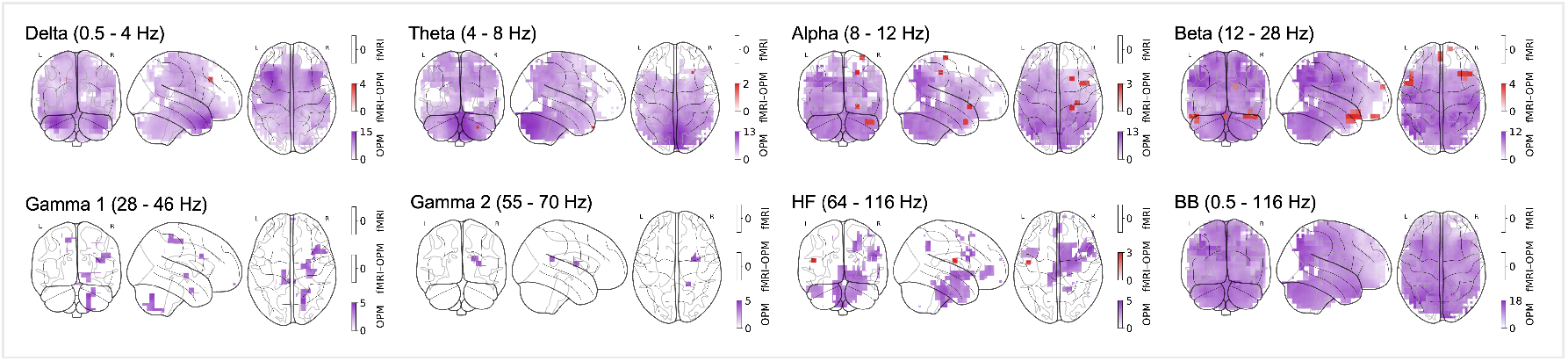
Reliable *z*-scores exclusive to between-subject and between-method comparisons for OPM and fMRI. Channels surviving MC correction for between-subject OPM but not between-subject fMRI (purple), between-subject fMRI but not between-subject OPM (note, no channels survived correction for MC in between-subject fMRI only), and between-method OPM-fMRI but not between-subject OPM nor fMRI (red). Color bars reflect *z*-scores (see above).

## Notes

### Competing Interest Statement

The authors have declared no competing interest.

### Summary of Updates

This version reflects a substantial revision of the manuscript; all primary analyses have been recomputed; all associated results, figures, tables, and Supplemental materials have been updated accordingly; new analyses further evaluating reliability and cross-modal comparisons are now reported in the Supplemental materials; the Discussion section has been expanded to address these revised and new results.

